# Whole-genome resequencing reveals exceptions to panmixia in the exploited top fish predator Antarctic toothfish (*Dissostichus mawsoni*)

**DOI:** 10.64898/2025.12.01.691552

**Authors:** Jilda Alicia Caccavo, Larissa S. Arantes, Enrique Celemín, Susan Mbedi, Sarah Sparmann, Camila J. Mazzoni

## Abstract

Antarctic toothfish (*Dissostichus mawsoni*) are a critical top fish predator in the Southern Ocean for whom a changing climate and fishing pressure challenge our understanding of their population dynamics. Their complex, long-lived life history presents challenges to understanding their population dynamics that we endeavor to overcome through the use of both reduced-representation sequencing and whole-genome resequencing data from fish across different cohorts. Our unique sample set allowed us to compare genomic structure, diversity, and demographic changes between cohorts born before and after the year 2000, when changes to fisheries management practices in the Southern Ocean reduced incidences of illegal, unreported, and unregulated fishing. While our population structure analyses corroborate recent studies showing panmixia across circumpolar populations of *D. mawsoni*, comparisons between cohorts revealed key exceptions. A lack of admixture in fish born before 2000 contrasted with widespread admixture among post-2000 individuals, consistent with increased mixing and/or genotype influx. Concernedly, genome-wide diversity declined over time in the Ross Sea (management area 88), with the significant losses observed in heterozygosity between pre-2000 and post-2000 cohorts. Elevated levels of inbreeding in the Ross Sea, coupled with evidence for recent population contraction from positive Tajima’s D values, provided potential explanations for the observed reductions in genomic diversity. Our results reconcile circumpolar panmixia with localized erosion of diversity and elevated inbreeding in the Ross Sea sector, underscoring the value of temporal monitoring and the need to prioritize future research on the interplay between fisheries pressure and environmental change in driving population dynamics and genomic structure in tootfhish.

## 1 Introduction

Marine species that are pelagic at any point during their life cycle often benefit from the connectivity afforded by oceanic currents to maintain high levels of gene flow between populations, even across large distances (Hellberg, 2009; Grummer et al., 2019; White et al., 2010; Schraidt et al., 2023; Liczner et al., 2024; Jahnke and Jonsson, 2022; Silva et al., 2019; Truelove et al., 2016). High levels of gene flow promote genetic diversity within and among populations, promoting population health and resilience to environmental and anthropogenic stressors (Vásquez et al., 2023; Hughes et al., 2008; Wernberg et al., 2018; Salo and Gustafsson, 2016). This phenomenon is particularly pronounced among fish species in the Southern Ocean, where multiple current systems promote genetic connectivity across the vast array of Antarctic fish species with pelagic periods during their life histories (Bernal-Durán et al., 2024; Caccavo et al., 2021; Matschiner et al., 2009; Damerau et al., 2012; Van de Putte et al., 2012). Exceptions to the high levels of genetic connectivity observed across Antarctic fish species have been linked to changes in current systems (Damerau et al., 2014; Agostini et al., 2015; Caccavo et al., 2018), environmental change (Desvignes et al., 2025), and overexploitation (Young et al., 2018). Species subject to all three phenomena are particularly vulnerable to disruptions in gene flow, resulting in a loss of genetic diversity, which can ultimately lead to extirpation, endangerment, and ultimately extinction (Pelletier and Coltman, 2018; Pinsky and Palumbi, 2014; Bijlsma and Loeschcke, 2012; Bosse and van Loon, 2022). Antarctic toothfish (*Dissostichus mawsoni*) are an example of one such species, because they: 1) rely on Southern Ocean current systems to complete their life cycle and exchange individuals between regional populations (Ashford et al., 2017; Hanchet et al., 2015a); 2) depend on sea ice, which has experienced record-breaking fluctuations and reductions across the Southern Ocean in recent decades (Josey et al., 2024; Eayrs et al., 2019; Liu et al., 2023; Turner et al., 2022), during their early life development (Parker et al., 2021, 2019); and 3) are subject to fisheries exploitation (Hanchet et al., 2015b; Abrams, 2014).

A top fish predator, *D. mawsoni* can reach up to 2 m in length, with a lifespan of up to 30 years, only reaching sexual maturity at 12 - 15 years of age (Hanchet et al., 2015a; Caccavo et al., 2021). This long-lived and slow-developing species has a complex life cycle (Figure 1): 1) benthopelagic adults spawn over offshore sea mounts (Hanchet et al., 2008); 2) positively buoyant eggs rise to the surface, where larvae hatch in the platelet ice environment beneath sea ice (Parker et al., 2021); 3) developing larvae are transported to the continental shelf via current systems linking offshore sea mounts to the continental slope (Behrens et al., 2021); 4) juveniles on the shelf gain condition on nutrient-rich prey (La Mesa et al., 2019; Hanchet et al., 2008); before 5) maturing into neutrally buoyant adults (Near et al., 2003; Fenaughty et al., 2008); that 6) ultimately repeat the cycle by following current systems that transport individuals from feeding areas on the continental slope to offshore sea mounts where spawning occurs (Di Blasi et al., 2024; Parker et al., 2019).

**Figure 1.**
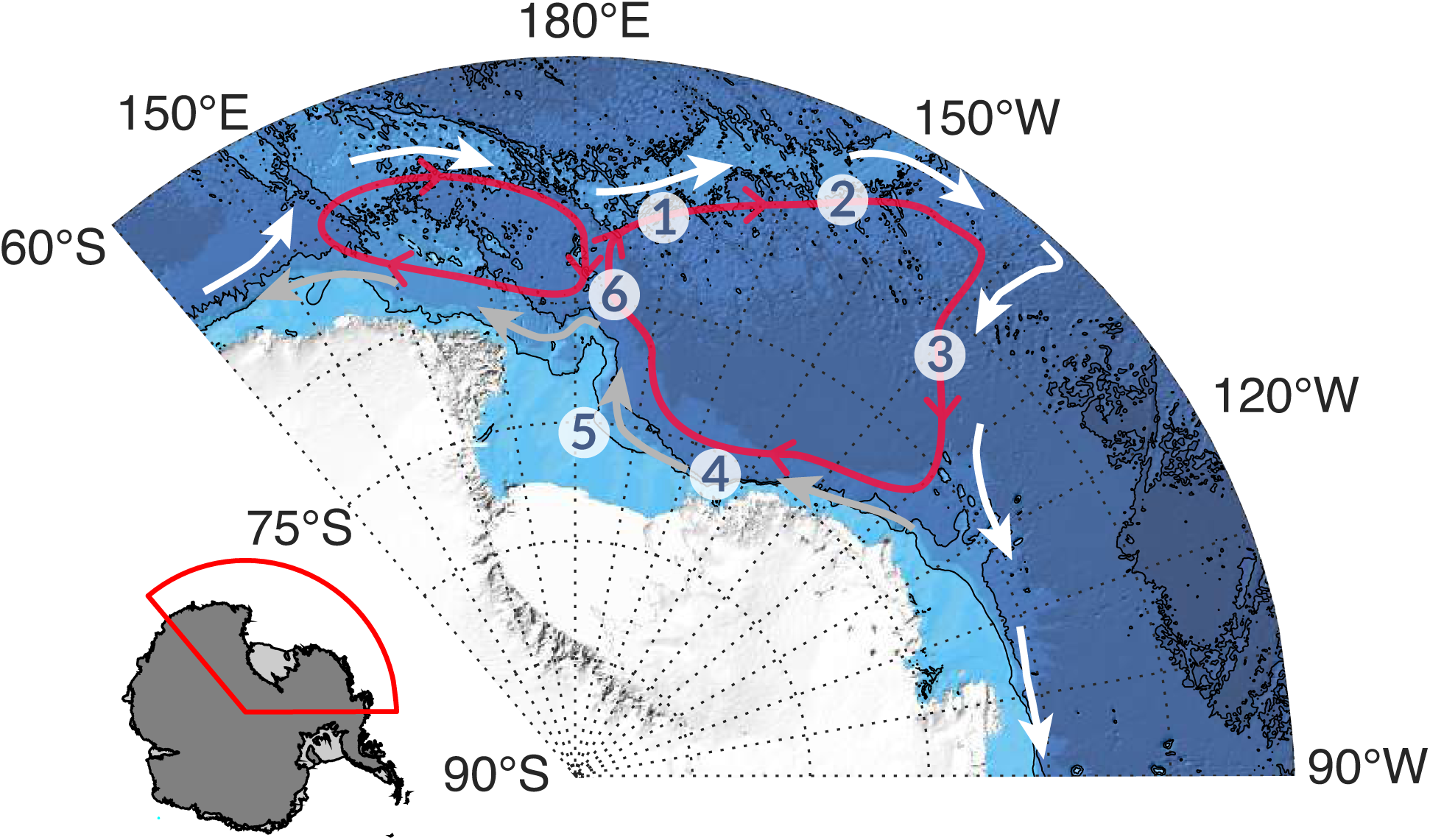
*D. mawsoni* life cycle, depicting: 1) offshore spawning over sea mounts, where 2) pelagic eggs and larvae develop near the surface under the sea ice, before 3) being transported to the continental shelf where 4) benthopelagic juveniles have access to nutrient-rich prey, and where 5) the accumulation of lipid stores during maturation permits adults to gain neutral buoyancy, before ultimately 6) repeating the cycle upon sexual maturity by following current systems from the shelf to offshore sea mounts where spawning occurs. While the life cycle is shown here in the Ross Sea (CCAMLR management area 88), it can be extrapolated to other circumpolar populations as well. White arrows depict the Antarctic Circumpolar Current (ACC) (Orsi et al., 1995), while the gray arrows indicate the relative position of the Antarctic Slope Current (ASC) along the continental slope at the 1000-m isoline. The red circles represent the Ross Gyre formed by the confluence of the ACC and ASC. Bathymetry data were pulled from IBSCO-v2 (Dorschel et al., 2022).

In addition to being an important prey species for apex predators such as whales and seals (Foster-Dyer et al., 2024; Rumolo et al., 2020; Ainley et al., 2020; Queirós et al., 2025; Pinkerton et al., 2010; Pinkerton and Bradford-Grieve, 2014; Bestley et al., 2020; Ainley and Ballard, 2012), *D. mawsoni* are of economic importance in that, together with the closely related Patagonian toothfish (*Dissostichus eleginoides*), they support the most lucrative fishery in the Southern Ocean, worth over 200 million USD annually (Grilly et al., 2015; Secretariat, 2018, 2019, 2022; Brooks, 2013). Marketed globally as “Chilean sea bass” (Mayyasi, 2014), interest in fishing for *D. mawsoni* began in the 1980s, before ramping up in the 1990s, a period characterized by illegal, unreported, and unregulated (IUU) fishing of *D. mawsoni* (Österblom and Sumaila, 2011; Lack, 2008; Ainley et al., 2012; Brooks, 2013), which led to recommended consumption bans by non-governmental organizations such as Seafood Watch (Cascorbi, 2006). Beginning in the early 2000s, the Commission for the Conservation of Antarctic Marine Living Resources (CCAMLR) implemented catch documentation and surveillance systems that reduced IUU fishing and improved the sustainable management of the fishery (Agnew, 2000; Miller, 2004; Kock, 2007; Baird, 2024). Starting in the Ross Sea region of the Southern Ocean (CCAMLR management area 88), the legal fishery expanded over time to include parts of East Antarctica (CCAMLR management area 58), as well as the Weddell Sea (CCAMLR management area 48) (Kock et al., 2007) (Figure 2b). Since these measures were introduced in the early 2000s, documented catches for *D. mawsoni* steadily increased in the following years before leveling off to the quantities observed today (CCAMLR, 2025b) (Figure 2c).

**Figure 2.**
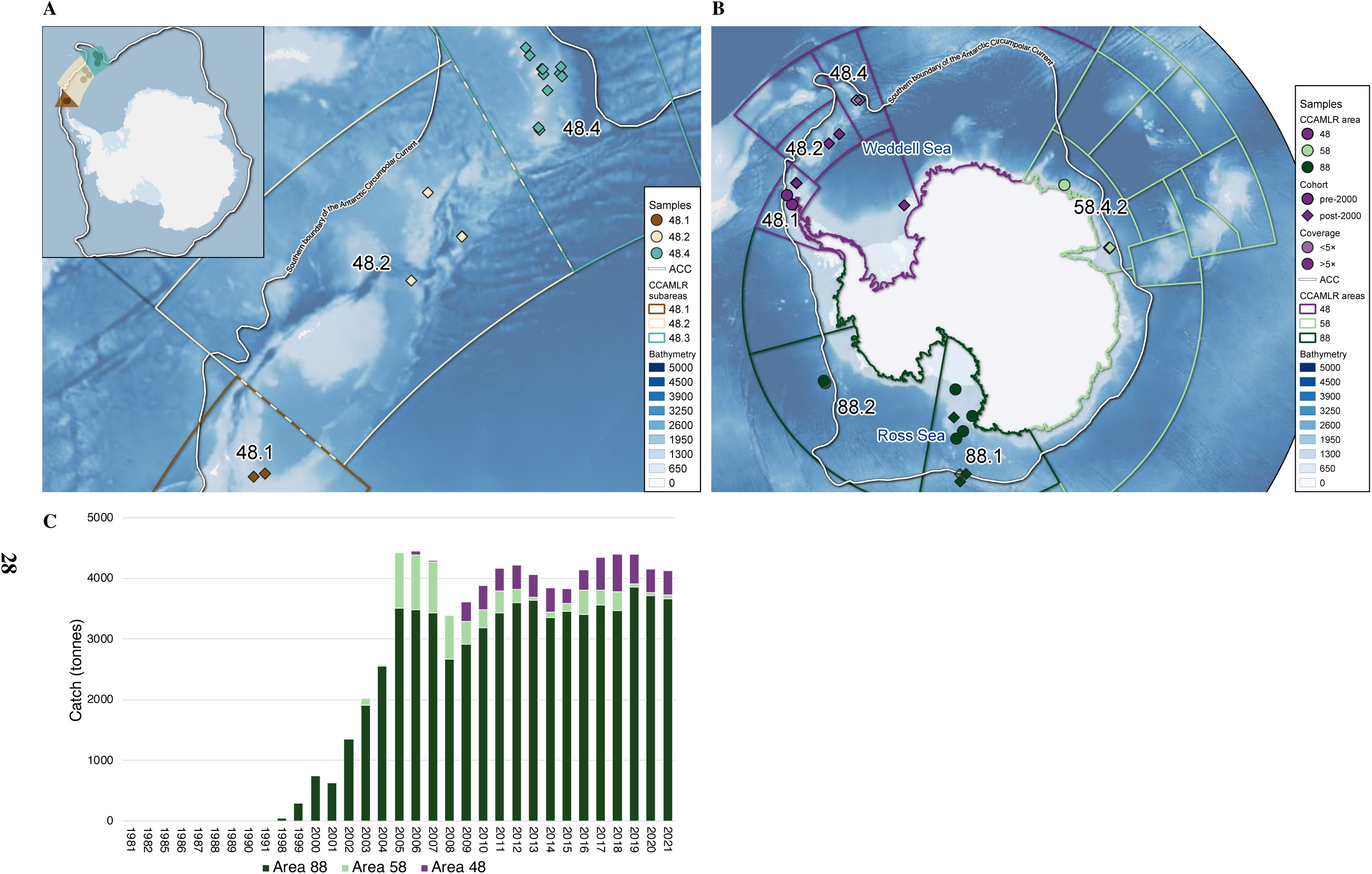
Maps showing the provenance of *D. mawsoni* samples from the (A) 3RADseq and (B) WGR datasets. Sample locations are presented based on various parameters detailed in the map legends. The Antarctic Circumpolar Current (ACC) is shown in white on both maps based on Orsi et al. (1995). CCAMLR management subareas and areas derive from CCAMLR (2017); relevant subareas and place names referred to in the text are labeled. Bathymetry data are based on IBSCO-v2 (Dorschel et al., 2022). Maps created with Quantarctica v.3.2 (Matsuoka et al., 2021) using QGIS v.3.16.7 (QGIS Development Team, 2023). (C) Catch in tonnes of *D. mawsoni* from 1981 to present (CCAMLR, 2025b).

Since the emergence of legal fishing for *D. mawsoni*, multiple studies have endeavored to understand its population structure through analysis of age-class distributions (Yates et al., 2019; La Mesa et al., 2019), tag recapture programs (Grilly et al., 2022), otolith chemistry (Carimán et al., 2025; Zhao et al., 2024; Ashford et al., 2012), and genetics (Parker et al., 2002; Smith and Gaffney, 2005; Kuhn and Gaffney, 2008; Mugue et al., 2014; Ceballos et al., 2021; Choi et al., 2021; Maschette et al., 2023). Early genetics studies yielded mixed results using low numbers (i.e., *n* = 10 markers) of various marker types. While Parker et al. (2002) identified significant differences between *D. mawsoni* populations in management areas 48 and 88 using randomly amplified polymorphic DNA, Smith and Gaffney (2005) found no differences in genetic variation among all three management areas (48, 58, and 88) based on markers derived from mitochondrial DNA and nuclear introns. Likewise, Kuhn and Gaffney (2008) reported genetic differentiation across all management areas of *D. mawsoni* using gene fragments and mitochondrial DNA markers. However, this finding was later challenged by Mugue et al. (2014), who, applying the same nuclear markers from Kuhn and Gaffney (2008) to a different cohort, observed genetic homogeneity across the species’ circumpolar distributions. With the increasing affordability of next-generation sequencing, recent studies have employed reduced-representation approaches (i.e., restriction site-associated DNA sequencing, RADseq), to test to test the null hypothesis that circumpolar distributions of *D. mawsoni* represent a single panmictic metapopulation throughout the Southern Ocean. RADseq analyses of fish from areas 48 and 88 (Ceballos et al., 2021), as well as across all circumpolar distributions (Maschette et al., 2023), failed to identify any evidence of restrictions to gene flow between regional populations of *D. mawsoni*, ultimately supporting the panmixia hypothesis. While microsatellite markers used in Choi et al. (2021) could not distinguish between fish from various parts of management areas 58 and 88, mitochondrial markers revealed significant differences between locations that increased proportionally with geographic distance.

The research into *D. mawsoni* genetic structure presents a paradox: if most studies support the hypothesis that gene flow homogenizes genetic differences across circumpolar distributions (Smith and Gaffney, 2005; Mugue et al., 2014; Ceballos et al., 2021; Maschette et al., 2023), why is it that there remains evidence across a handful of studies that regional populations can achieve levels of isolation sufficient to support significant differences in genetic composition (Parker et al., 2002; Kuhn and Gaffney, 2008; Choi et al., 2021)? The answer may ultimately lie in a revised hypothesis that proposes that both possibilities may be true at the same time. A key consideration absent from previous studies of *D. mawsoni* genetic structure is cohort. As has been observed generally in broadcast spawners (Vendrami et al., 2021), and particularly in Antarctic fish (Agostini et al., 2015; Caccavo et al., 2018; Papetti et al., 2007, 2012), genetic differentiation can vary over time between cohorts, as environmental or anthropogenic influences on recruitment (i.e., the proportion of fish reproducing in a given year) shift genetic diversity within and thus between populations over time, in a phenomenon known as chaotic genetic patchiness (Johnson and Black, 1982; Eldon et al., 2016). It is thus possible that connectivity between circumpolar distributions of *D. mawsoni* varies over time, resulting in a situation in which certain cohorts exhibit no genetic differences between regional populations, while others that have experienced significant interruptions in gene flow can become genetically isolated. Because genetically isolated populations can experience reductions in genetic diversity that render them less resilient to anthropogenic pressures and environmental change, precautionary fisheries management requires an understanding of how the genetic connectivity of an exploited species may change over time (Benestan, 2019).

To address this gap in our understanding of *D. mawsoni* genetic structure over time, essential to the continued sustainability of its fishery, we use both RADseq and whole-genome resequencing (WGR) approaches to investigate genetic structure on regional and circumpolar scales. In particular, we infer population structure and estimate levels of genetic diversity and effective population size across both management areas and between cohorts. While RADseq approaches can generate genome-wide markers of genetic diversity (singlenucleotide polymorphisms, SNPs) representative of approximately 1% of the genome, WGR approaches can create SNPs representative of >70% of the genome, producing on the order of 10^5^ to 10^6^ SNPs (Therkildsen and Palumbi, 2017; Fuentes-Pardo and Ruzzante, 2017). This greater representation of the genome permits the analysis of demographic trends over time, such as changes to effective population size, and population expansions and contractions (Martchenko and Shafer, 2023; Lowry et al., 2017). Thanks to recent work from our research group (Caccavo et al., 2024), in addition to the standard sample type of muscle tissue or fin clips preserved in alcohol, we are able to include in our study DNA extracted from trace tissue adhered to the surface of *D. mawsoni* otoliths, or ear bones. Because otoliths are collected every year by the fishery to analyze ageclass distributions (CCAMLR, 2025a), this sample type is available from across all circumpolar distributions of *D. mawsoni*, from the early years of the fishery in the beginning of the 2000s, through to the present (Welsford et al., 2009; Toomey et al., 2016). The ability to use otoliths to extract DNA, combined with access to some of the same samples used in Kuhn and Gaffney (2008), allows us to compare genetic structure and demographic trends over time between cohorts of fish exposed to greater levels of IUU fishing before the 2000s, and fish born after the 2000s, when improved management strategies by CCAMLR reduced incidences of IUU fishing.

## 2 Materials and Methods

### 2.1 Samples

Two sets of samples were used in this study (Figure 2). First, a set of 108 tissue (muscle and fin clip) samples from key CCAMLR management subareas within the Weddell Sea (subarea 48.1 *n* = 40, subarea 48.2 *n* = 41, subarea 48.4 *n* = 28) (Figure 2a). Second, a set of 41 samples consisting of both tissue and otolith samples from across all three CCAMLR management areas (area 48 *n* = 13, area 58 *n* = 8, area 88 *n* = 20) (Figure 2b). To produce the 108-sample set from the Weddell Sea, we employed a reduced-representation sequencing approach (3RADseq) to process 192 *D. mawsoni* samples in two 96-well plates. These 192 samples were drawn from 456 tissue samples generously provided by 4 countries (New Zealand, Ukraine, United Kingdom and United States). Samples were selected for inclusion in the preparation of 3RADseq libraries to maximize representation of CCAMLR management subareas across the Weddell (subareas 48.1 *n* = 48, 48.2 *n* = 48, 48.4 *n* = 58) and Ross (subarea 88.1 *n* = 38) Seas, while also standardizing cohort representation within subareas based on total fish length (TL) using age-length data from Horn (2002) and Brooks et al. (2011). Samples from different management subareas were then selected randomly for inclusion in 3RADseq plates, on which they were randomly distributed to avoid plate effects such as pipetting error from being restricted to certain geographic groups. The circumpolar 41-sample set consisted of lower quality DNA samples extracted from trace tissue remaining on the exterior of fish otoliths, as well as from poorly-preserved tissue samples, thus benefiting from a whole-genome resequencing approach to reduce artefacts and biases (Arantes et al., 2023; Caccavo et al., 2024; Schultz et al., 2022; Valière et al., 2007). The 41 *D. mawsoni* WGR libraries used in this study are the same as those presented in Caccavo et al. (2024), where extensive details on library preparation can be found.

### 2.2 RAD sequencing

The 108-tissue sample set was prepared using the 3RADseq method detailed in Arantes et al. (2023), which is a modified version of the protocol described in Bayona-Vásquez et al. (2019) to include a primer with unique molecular tags (Hoffberg et al., 2016). Reads from loci prepared via 3RADseq were assessed for coverage through smallscale sequencing (hereafter referred to as “spike-in”), which permitted re-pooling to achieve greater uniformity in read coverage across samples, thus requiring less high-output sequencing to achieve the desired minimum 30×-coverage across all samples (Arantes et al., 2023).

#### 2.2.1 Library construction

We began by simulating the digestion of the *D. mawsoni* reference genome (GenBank accession number JAAKFY000000000) with various combinations of different restriction enzymes, with the aim of achieving a set of approximately 25,000 loci. Based on this approach, we selected EcoRI, MspI and ClaI which allowed us to produce nearly 26,000 loci between 325 and 550 bp.

We added 200 ng of DNA to the digestion reaction, which included 10 units of each enzyme, NEB 10× CutSmart Buffer, and a unique combination of dual internal barcodes (5 *µ*M) for each sample, in a total volume of 15 *µ*L. After 2 h of digestion at 37°C, DNA ligase and ATP were added to the reaction and the temperature was cycled to encourage ligation (22°C) followed by digestion (37°C), with a final 20-minute cycle at 80°C. Following the digestion/ligation reaction, the 192 libraries were multiplexed into 4 pools of 48 samples each, and ligation products were cleaned using 0.8× CleanPCR magnetic beads (GC biotech). The final digestion/ligation product consisted of 20 *µ*L; we used 5 *µ*L of this product from each sample to create the 48-sample pool. Despite each reaction benefiting from the input of equal quantities of DNA, we nonetheless observed significant variability in read coverage after spike-in, which may be related to pipetting error and variable DNA fragmentation among samples, resulting in inconsistent patterns of digestion. Our spike-in strategy allowed us to avoid situations in which read coverage was so divergent across the pooled sample set that it would be impossible to adjust pooling volumes to normalize read coverage across the sample pool (Arantes et al., 2023).

Size selection was performed on cleaned sample pools using a BluePippin 1.5% cassette (Sage Science). Two size selections were performed - one at 430 - 580 bp and one at 580 - 730 bp. These bp ranges were chosen based on optimization trials of the protocol, which revealed that BluePippin size selection at 430 - 580 bp and 580 - 730 bp resulted in fragment ranges of 345 - 495 bp and 495 - 645 bp respectively. This discrepancy in fragment sizes is related to both the length of the adapter sequences (80 bp), as well as a tendency for our BluePippin device to tend towards narrower and shorter size ranges. While we only performed sequencing on the set of longer fragments (495 - 645 bp), we found that it was necessary to perform size selection on the shorter fragment sizes at the same time to ensure the removal of fragments <495 bp from the fragment pool.

A single-cycle PCR was then performed to incorporate iTru5-8N primers into the size-selected fragments in four replicate reactions using 10 *µ*L each of the total 40 *µ*L BluePippin elution. Afterwards, each replicate reaction was cleaned with CleanPCR magnetic beads in preparation for the indexing PCR, which added different external P7 and P5 indexes, permitting libraries to be combined for sequencing. The number of cycles of this final indexing PCR was determined based on the amount of input DNA; ultimately we found that 10 cycles allowed for the best compromise between adequate resultant DNA concentrations without excessive PCR duplicates. In addition, we performed the extension step (72°C) of the PCR for 1 minute instead of 30 seconds to accommodate larger fragment sizes (i.e., 495 - 645 bp). At this stage, all replicate reactions were recombined into a single pool, equivalent to the original maximum 48-sample pool created after digestion/ligation. After a final cleaning step with CleanPCR magnetic beads, we used High Sensitivity DNA ScreenTape on an Agilent Tapestation, to verify fragment size distributions and DNA concentration of our final 3RADseq library pool prior to spike-in.

#### 2.2.2 Preprocessing

We followed the spike-in strategy detailed in Arantes et al. (2023), performing bioinformatics preprocessing of reads using the Snakemake workflow, which included: 1) phiX control library cleaning; 2) demultiplexing; 3) filtering of PCR duplicates; 4) concatenating of the four replicates of each library; 5) merging of forward and reverse reads; and 6) a final quality filtering of the reads. For our study, this spike-in strategy served two key purposes: 1) to check the balance of individuals within the library pool based on read coverage; and 2) to confirm the overlap of fragment length distributions in the library pool, which assures that the same loci are represented across individuals. After analysis of the initial spike-in data, 42/48, 13/48, 39/48, and 15/48 libraries respectively were retained for equimolar repooling to improve read balance. The tissue quality of the 38 Ross Sea samples was relatively poor, resulting in none of the libraries producing enough reads to be included in library pools. For this reason, tissue samples from the Ross Sea were incorporated into the WGR dataset. All aforementioned library preparation steps were then repeated on the balanced library pool. Spike-in sequencing was then performed on the final 3RADseq library pools to confirm balance before combining all 3RADseq libraries with all whole-genome resequencing (WGR) libraries (see section 2.3 Whole-genome resequencing) to allow for increased sequencing diversity across cycles. A final spike-in assessment on the combined 3RADseq- and WGR-library pool was performed prior to 150-bp paired-end high-output sequencing at the Competence Centre for Genomic Analysis (CCGA) Kiel, Germany, using one lane of an Illumina NovaSeq S4 flow cell.

#### 2.2.3 SNP calling

Preprocessing of high-output 3RADseq reads was performed in the same manner as 3RADseq spike-in data, with the following modifications: 1) no phiX control library cleaning was needed, as no phiX was added to the high-output sequencing run; 2) merging was not required, as reads were only 150 bp, compared to the 300-bp reads sequenced during spike-in; 3) Cutadapt (Martin, 2011) was used to trim adapters from 3’-sequence ends; 4) chimeric and undigested sequences were removed; and 5) reads were mapped to the *D. mawsoni* reference genome (GenBank accession number JAAKFY000000000).To identify SNPs, we used the STACKS reference-based pipeline v.2.61 (Catchen et al., 2011; Rochette and Catchen, 2017) calling variant sites at loci genotyped in at least 60% of all individuals sequenced (*n* = 109), and performing population-level filters using parameters *p* = 1 and *r* = 0.6. Further filtering was performed using VCFTOOLS v.0.1.16 (Danecek et al., 2021), removing sites with a coverage of <10× (`--min-meanDP 10`, `--min-DP 10`) or >120× (`--min-meanDP 120`, `--min-DP 120`), with >25% missing data (`--max-missing 0.75`), or linked SNPs (`--thin 500`). To assure that none of the 109 individuals in our dataset were first-degree relatives, we used the `--king-cutoff` option in PLINK2 v.v2.00a3.7LM (Chang et al., 2015) to perform relation-based pruning. This resulted in the removal of one sample, leaving 108 individuals in the final dataset. Coverage of 3RADseq libraries ranged from 23.03× - 46.68× (average = 32.63×) after all filtering steps.

### 2.3 Whole-genome resequencing

The circumpolar 41-sample set is composed of the same samples presented in Caccavo et al. (2024), where extensive details can be found on DNA extraction, whole-genome resequencing (WGR) library preparation, sequencing, bioinformatics processing, and SNP calling.

#### 2.3.1 Library construction

In short, otolith DNA was extracted from trace tissue remaining on the surface of otoliths using the E.Z.N.A® MicroElute® Genomic DNA Kit (omega bio-tek), based on an optimized version of the manufacturer’s protocol. DNA from poorly-preserved tissue samples was extracted using the Qiagen DNeasy blood & tissue kit following the manufacturer’s instructions. Whole-genome libraries from both otolith and tissue DNA extracts were prepared using NEBNext® Ultra™ II FS DNA Library Prep Kit for Illumina® based on an optimized version of the manufacturer’s protocol. Whole-genome libraries were selected in line with the strategy outlined in Arantes et al. (2023), which used spike-in to select libraries which met minimum mapping and deduplication criteria. Selected libraries were then combined with 3RADseq libraries (see section 2.2 RAD sequencing) for high-output sequencing.

#### 2.3.2 Preprocessing and SNP calling

Bioinformatics preprocessing was performed on the raw NovaSeq reads to demultiplex samples, map them to the *D. mawsoni* reference genome, and remove PCR duplicates. Then, reads were filtered further for the presence of indels, coverage, and mapping and base quality. When checking the final coverage of our preprocessed WGR libraries using SAMTOOLS v.1.20 (Danecek et al., 2021), we found that there was a wide range of coverage levels across libraries (1.13× – 27.09×). Given the potential impact that differing coverage levels can have on SNP calling (e.g., heterozygosity bias) (O’Leary et al., 2018), we opted to create three datasets downsampled to different coverage levels: 1) downsampled to 2× (*n* = 37); 2) downsampled to 5× (*n* = 24); and 3) not downsampled (*n* = 41) (Table S1). This allowed us to compare the impact of coverage levels on subsequent SNP calling and downstream analyses. Downsampling was performed using SAMTOOLS `view`, with the downsampling parameter `-s` set to the proportion of reads to retain in order to achieve the desired coverage. After checking the coverage post-downsampling, the downsampling parameter was tweaked iteratively to attain a final coverage within *±*0.05× of the target downsampled coverage. Note that among neither the 3RADseq nor WGR datasets were sex-linked scaffolds filtered, because *D. mawsoni* do not have heterochromatic sex chromosomes (Ghigliotti et al., 2007, 2016).

Two approaches were used to call SNPs: 1) genotype likelihoods, which account for the uncertainty inherent to low-coverage data (Lou et al., 2021); and 2) genotypes, which are necessary for certain analyses of demographic history. Genotype likelihoods were derived using ANGSD v.0.933 (Korneliussen et al., 2014), while to call genotypes, we used BCFTOOLS v1.17 (Danecek et al., 2021). For downstream analyses that required unlinked SNPs, we used NGSLD v.1.2.0 (Fox et al., 2019) and PRUNE GRAPH to identify and filter linked SNPs respectively from genotype likelihoods, while for genotypes, we used PLINK2 `---indeppairwise`. For both genotype likelihoods and genotypes, we applied a 20 kb cutoff to identify linked SNPs, using a minimum weight setting of 0.5.

### 2.4 Population analyses

#### 2.4.1 Genomic summary statistics

We estimated genome-wide heterozygosity among WGR libraries in ANGSD based on the folded site frequency spectrum (SFS). To derive the folded SFS, we used the reference genome as the ancestral state to first calculate the folded site allele frequency likelihood, before then computing the folded SFS. Based on the same SFS, we also derived Watterson’s theta and Tajima’s D to consider population mutation rate and neutral versus non-random selection processes respectively. These were calculated using a sliding window approach, with window size set to 50 kb at a step size of 10 kb. Finally, we derived individual inbreeding coefficients (*F*) using NGSF v.1.2.0 (Vieira et al., 2013). We first ran NGSF to attain an approximate *F* value per individual, using the `--approx_EM` method with a maximum root mean squared difference between iterations (`---min_epsilon`) set to 1×10^−5^, starting with random values. The output of this initial run was used to set starting parameters for the final run, with `--min_epsilon` decreased to 1×10^−7^. This two-step process was replicated 10 times to avoid convergence to local maxima (Vieira et al., 2013). Differences in population statistics between geographic and cohort groups were assessed in base R using parametric one-way analysis of variance (ANOVA), followed by post hoc *t*-test with Holm correction.

#### 2.4.2 Neutral and adaptive diversity

Adaptive processes have the potential to shape genetic structure, and thus must be accounted for in analyses of population structure (Chung et al., 2023). To this end, we performed outlier SNP analyses on both 3RADseq and WGR datasets. For the 3RADseq dataset, we used 4 approaches to identify outlier SNPs: 1) PCADAPT, which uses principal component analysis (PCA) to define outliers based on population structure (Luu et al., 2017). We used K = 2 principal components because this was the K value that minimized the genomic inflation factor as derived from the scree plot (Figure S1a); 2) BAYESCAN, which applies a Bayesian approach to identify outliers based on their probability of deviating from neutral evolutionary models (Foll and Gaggiotti, 2008); 3) OUTFLANK, which relies on a likelilhood approach to determine the null distribution of population differentiation for neutral loci without taking demographic history into consideration (Whitlock and Lotterhos, 2015); and 4) FDIST implemented in ARLEQUIN, which identifies loci with elevated *F*_ST_ for simulated background heterozygosity under island and hierarchical island models (Beaumont and Nichols, 1996). For the WGR dataset, we used 3 approaches to identify outlier SNPs: 1) PCADAPT; 2) BAYESCAN; and 3) OUTFLANK. For PCADAPT, we used K = 2 principal components based on the scree plot of the WGR data, which showed that genomic inflation factor was minimized at this value (Figure S1b). All analyses were run with default parameters.

#### 2.4.3 Population structure

We analyzed population structure on unlinked SNPs derived from both the 3RADseq and WGR libraries using admixture and principal component analysis (PCA). For 3RADseq SNPs, we used Admixture (Alexander et al., 2009) to assign probabilities of individuals to genetic clusters K = 2 - 10, with each K value evaluated in parallel over 5 replicates. Admixture outputs were plotted using a dedicated R script (see article GitHubrepository). The bestfit K value maximizing classification was selected as K = 3 based on cross-validation. We performed PCA on 3RADseq SNPs using PLINK2 `--pca`. For WGR libraries, admixture and PCA were performed on genotype likelihoods using NGSADMIX (Skotte et al., 2013) and PCANGSD (Meisner and Albrechtsen, 2018) respectively. We assessed convergence in the admixture analysis by performing 20 independent runs, varying K between 2 and 11, at a minimum tolerance of 1×10^−10^ and a minimum likelihood ratio of 1×10^−6^. As with the 3RADseq dataset, K = 3 was selected as the bestfit K value for the WGR dataset.

### 2.5 Demographic history

#### 2.5.1 Runs of homozygosity

To assess genome-wide runs of homozygosity (RoH), we used PLINK2, with parameters based on those defined in Meyermans et al. (2020): 1) minimum number of SNPs per RoH (`--homozyg-snp 50`); 2) length in kb of the sliding window (i.e., the minimum RoH size, `--homozyg-kb 100`); 3) maximum distance in Mb between two adjacent SNPs to be considered as part of different segments (`--homozyg-gap 1000`); 4) number of SNPs a sliding window must have (i.e., the window size in number of SNPs, `--homozyg-window-snp 50`); 5) maximum number of heterozygous SNPs per window (i.e., the number of heterozygous SNPs allowed in a window, `--homozyg-window-het 4`); 6) number of missing calls allowed in a window (`--homozyg-window-missing 5`); 7) proportion of overlapping windows that must be called homozygous to define a given SNP as being in a “homozygous” segment (`--homozyg-window-threshold 0.05`); and 8) minimum density of SNPs to call an RoH (i.e., 1 SNP per how many kb, `--homozyg-density 100`). This analysis was run on the WGR SNP set based on genotypes. To balance the need for higher coverage to call genotypes with the need for adequate sample sizes among groups, we only analyzed RoH in the 5× dataset. To calculate the inbreeding level (F_ROH_), we divided the length in Mb of a given RoH by the size of the *D. mawsoni* genome (924.7 Mb).

#### 2.5.2 Effective population size (*N*_e_)

We used SMC++ v.15.5 to investigate changes in effective population size (*N*_e_) over time (Terhorst et al., 2017). As with the analysis of RoH, we restricted our investigation of *N*_e_ to called genotypes from the WGR dataset downsampled to 5×. First, we restricted our analysis to SNPs from scaffolds larger than 1Mba (28 scaffolds). Then we filtered SNPs that were not in Hardy–Weinberg Equilibrium (`--hwe 0.01`), followed by filtering SNPs that did not meet minimum base pair (`---QUAL<20`) and mapping (`--MQ<30`) quality criteria. In addition, we retained only SNPs present in at least 75% of individuals (`--AN/2<0.75`*n*) that were biallelic (`--max-alleles 2`), excluding indels (`--exclude types indels`). Notably, for analysis of *N*_e_, we did not filter for minor allele frequency, as the full allele frequency spectrum is necessary to effectively identify ancient bottlenecks or population contractions, as well as to reduce bias towards more recent demographic events (Terhorst et al., 2017). As suggested by Terhorst et al. (2017), we created composite likelihoods by varying the distinct individual (`-d`) in each of the three management areas. Four distinct individuals were chosen from each area, as *n* = 4 was the minimum sample size among management area groups. Distinct individuals were chosen to maximize coverage and representation of 3 different sample parameters: 1) sample type; 2) cohort; and 3) subarea. We calculated population size histories with the option `estimate`, using a generation time of 15 years (Parker and Grimes, 2010; Hanchet et al., 2015a; Caccavo et al., 2021) based on age-at-spawning (Kim et al., 2019; Lu et al., 2022), and a mutation rate of 2.85 × 10^−8^, as defined for *Chaenocephalus aceratus* when looking at *N*_e_ using SMC++ (Lu et al., 2022). While a lower mutation rate of 3.28 × 10^−9^ was used with PSMC (Kim et al., 2019) on another closely-related notothenioid, *Chionodraco hamatus*, we opted to use the higher mutation rate of 2.85 × 10^−8^ from Lu et al. (2022) because they employ the same analysis approach in SMC++. Based on Near et al. (2018), there is no phylogenetic argument for choosing the mutation rate for *C. aceratus* over *C. hamatus*, as both notothenioids are similarly related to *D.mawsoni*. All other parameters in the calculation of population size histories were kept as default (`---timepoints 33 100000 -c 50000 -rp .1 --knots 60 --spline cubic`). To visualize *N*_e_ over time by management area, we used the option `plot` to export the *json* files produced from the *N*_e_ calculations by population into a *csv* file, which was then used as input to create plots by mutation rate (2.85×10^−8^ and 3.28×10^−9^) with a dedicated R script (see article GitHub repository).

## 3 Results

### 3.1 SNP calling

After calling variant sites, filtering, and removing related individuals (1/109), we ended up with a set of 19,894 SNPs across 108 3RADseq libraries. For the 41 WGR libraries, we produced 1.9 - 5.4 million filtered SNPs across the 3 datasets based on genotype-likelihoods, with 0.8 - 2.8 million unlinked SNPs remaining after linkage disequilibrium pruning (Table 1). No outlier SNPs were identified within the 3RADseq dataset across all approaches. Because we could not be confident in called genotypes below 5× coverage, we only used the 5× dataset to call SNPs based on genotypes, which resulted in 2.8 million filtered SNPs, of which 2.4 million were unlinked (Table 1). While BAYESCAN and OUTFLANK failed to detect any outlier SNPs among unlinked SNPs derived from WGR, PCADAPT identified over 121 thousand putatively adaptive SNPs (Table 1). All subsequent analyses using genotypes were based on the SNP set from which all putatively adaptive SNPs were removed. Both to maintain consistency between the analyses run based on genotypes, as well as to maximize sample inclusion while also maximizing coverage, we present the results of analyses based on genotype-likelihoods from the 5× dataset. Replications of these same analyses in the 2× and noDS datsets are presented as Supplementary figures. Group sizes (*n*) in the 3RADseq and WGR datasets within management subareas and areas, as well as across cohorts, are summarized in (Table S1).

**Table 1.**
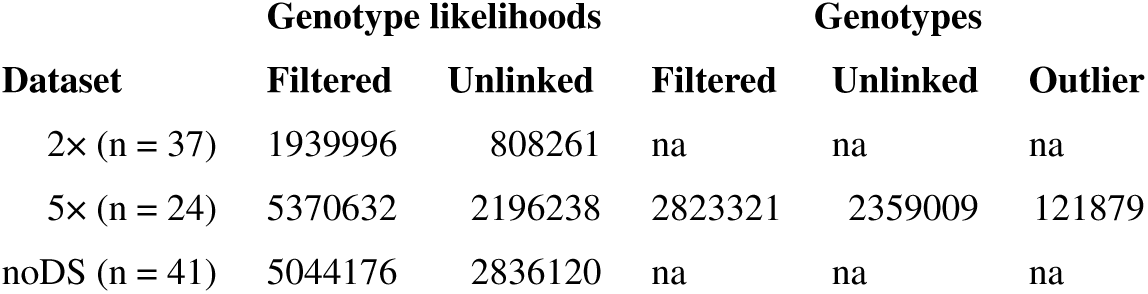
Total number of whole-genome library SNPs by dataset. Filtered SNPs have been treated for all SNP filters. Unlinked SNPs are filtered SNPs for which linkage disequilibrium pruning has removed linked SNPs. Outlier SNPs are filtered, unlinked SNPs identified as putatively adaptive SNPs in outlier scans. 2×, downsampled to 2×; 5×, downsampled to 5×; noDS, not downsampled; na, not applicable.

| Dataset | Genotype likelihoods |  | Genotypes |  |  |
| --- | --- | --- | --- | --- | --- |
|  | Filtered | Unlinked | Filtered | Unlinked | Outlier |
| 2× (n = 37) | 1939996 | 808261 | na | na | na |
| 5× (n = 24) | 5370632 | 2196238 | 2823321 | 2359009 | 121879 |
| noDS (n = 41) | 5044176 | 2836120 | na | na | na |

### 3.2 Genomic structure

In the Weddell Sea, there was no evidence of genomic structure based on SNPs derived from 3RADseq, neither from PCA or admixture analysis (Figure 3). In the PCA, the first principal component explained 10.2% of the variance, and created one outlier, sample 282 (Figure 3a). The second principal component also explained 10.2% of the variance, and created a second outlier, sample 309 (Figure 3a). Beyond these two outliers, the PCA showed no indication of clustering, consistent with high enough levels of gene flow between the Weddell Sea management subareas to consider them as one genetic population. When re-running the PCA with the outlier samples 282 and 309 removed (*n* = 106), no patterns of clustering emerged, though three additional samples (204, 274 and 212) were distinguished from the main sample cluster by the first principal component (ex-plaining 10.2% of the variance), while the second principal component (explaining 10.1% of the variance), separated outlier samples 204 and 274 from outlier sample 212 (Figure S2). Re-running the PCA after removal of these additional 3 outliers (*n* = 103) did not produce further outliers, nor did it reveal any previously unseen genetic structure, with both principal components explaining similar proportions of the variance (10.2% for both the first and second components) (Figure S2). Results from the admixture analysis showed a persistent lack of population structure across the Weddell Sea, as seen from the PCAs (Figure 3b).

**Figure 3.**
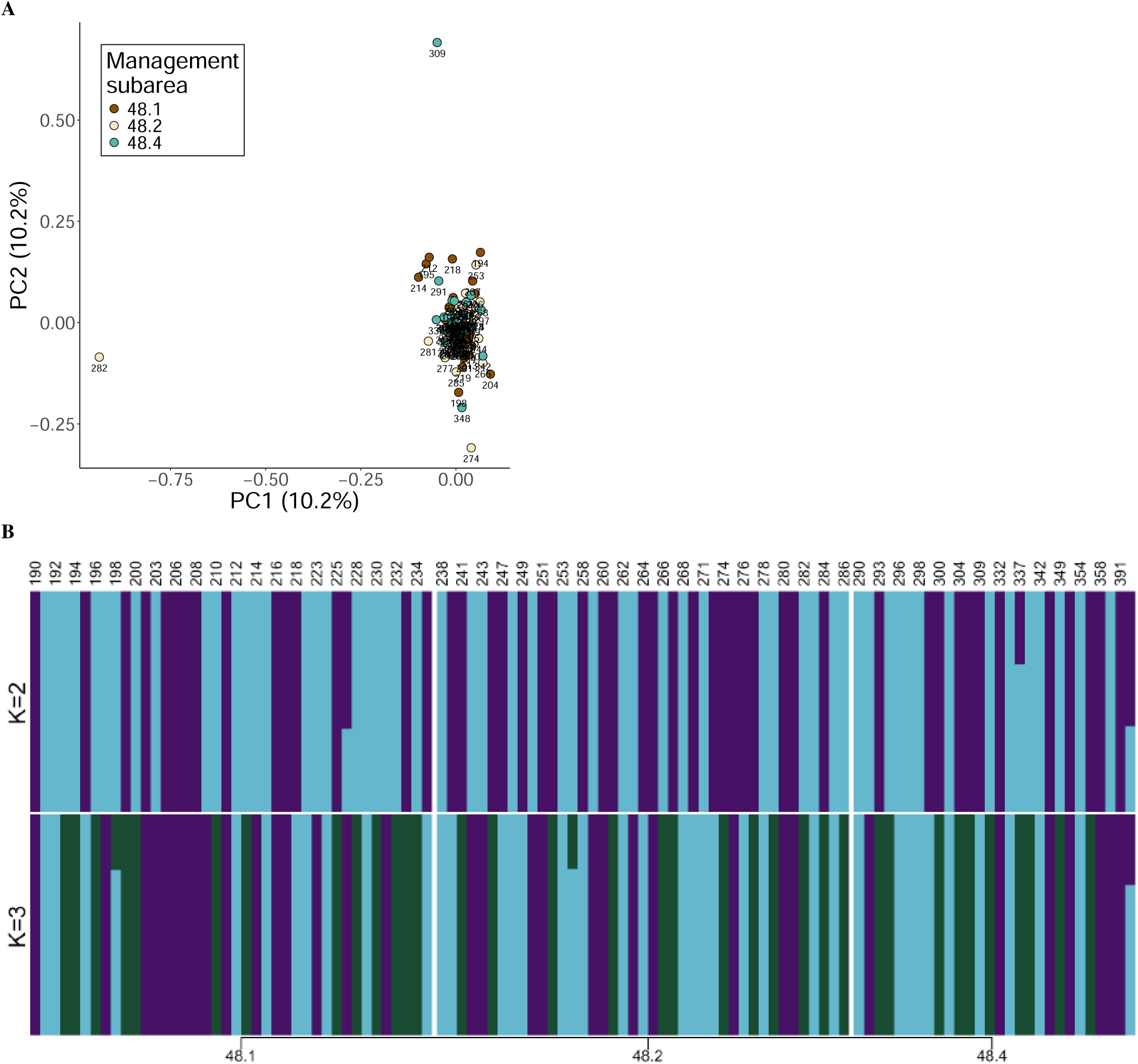
Population structure of *D. mawsoni* in the Weddell Sea based on 3RADseq data (*n* = 108). Groups labeled by CCAMLR management subarea (48.1, 48.2, 48.3). (A) PCA based on all samples. (B) Admixture plot of population structure for K = 2 and K = 3, with sample labels listed above the plot. Colors refer to admixture proportions and are not indicative of management subareas.

Geographic population structure remains absent on a circumpolar scale based on genotype likelihoods derived from the WGR dataset, though groupings based on cohort may reveal underlying patterns of change in population structure over time (Fig-ure 4). In the PCA, the first principal component explained 6.4% of the variance, while the second principal component explained 4.8% of the variance (Figure 4a). While 7 samples fell outside of the main PCA cluster, we did not consider these as outliers per se, given their large number (>5) and stochastic positioning. Alternative groupings of samples based on coverage, cohort, or sex did not reveal any quality distinguishing these 7 samples from those in the main cluster (Figure S3). Admixture did not correspond to CCAMLR management area (Figure 4b), though a pattern began to emerge when grouping samples by cohort (Figure 4c). Because *D. mawsoni* growth rates slow following sexual maturity at approximately 15 years of age (Horn, 2002; Brooks et al., 2011; Hanchet et al., 2015a; Caccavo et al., 2021) we were only able to estimate birth years to a precision of five years based on total fish length (TL) at the time of capture (Table S2). Given the low number of pre-2000 fish generally across our datasets (Table S1), as well as the key turning point that the year 2000 represents with respect to improved *D. mawsoni* management practices and reduced incidence of IUU fishing (Agnew, 2000; Miller, 2004; Kock, 2007; Baird, 2024), we chose to group samples into two different cohorts: fish estimated to have been born before the year 2000 (pre-2000), and fish estimated to have been born after the year 2000 (post-2000) (Figure S4). When organized into cohort groups, it became evident that pre-2000 fish do not show evidence of admixture (with the exception of sample 9544), while post-2000 were all admixed (with the exception of sample 35-431) (Figure 4c). These same patterns of population structure were reflected in the 2× dataset for both PCA and admixture analyses, (Figure S5), and in the noDS dataset for the PCA analysis (Figure S6). We were unable to run admixture on the noDS dataset due to computing resource constraints given the high levels of coverage of certain individuals (*n* = 14 >10×, *n* = 2 >20×, Table S2).

**Figure 4.**
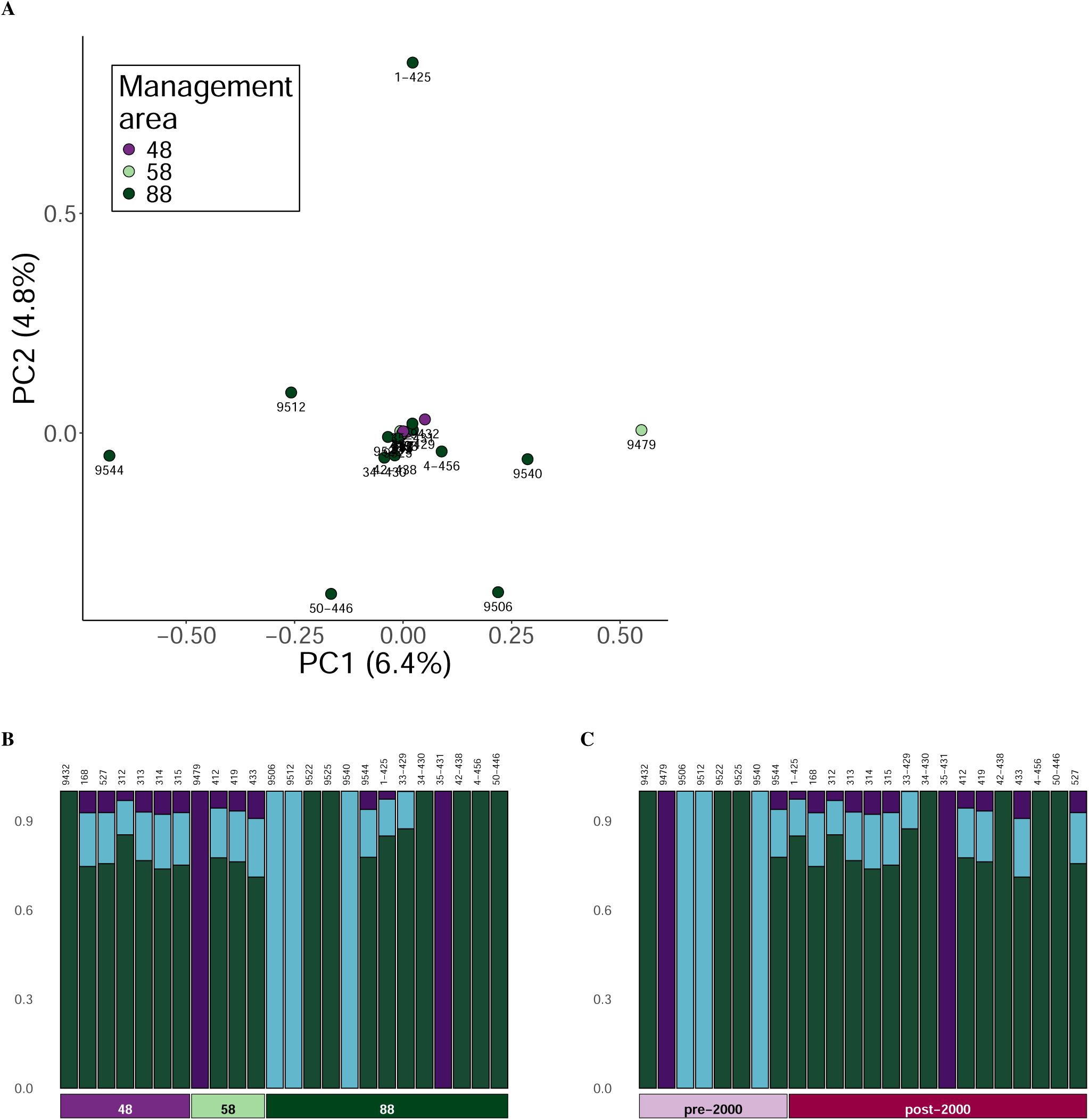
Circumpolar population structure of *D. mawsoni* based on WGR data (5× dataset, *n* = 24). Groups labeled by CCAMLR manage-ment areas (48, 58, 88). (A) PCA based on all samples, data labels indicate sample IDs. Admixture plots of population structure for *K* = 3, with sample IDs listed above the plot; ordered by (B) CCAMLR area. (C) cohort. Vertical bar colors refer to admixture proportions and are not indicative of management areas or cohorts.

### 3.3 Demographic history

Measures of heterozygosity revealed regional differences in genomic diversity within the Weddell Sea and across circumpo-lar toothfish distributions (Figure 5). Expected heterozygosity (*H_e_*) calculated based on 3RADseq data from the Weddell Sea (Figure 5a) indicated higher levels of diversity in CCAMLR management subarea 48.4 with respect to subareas 48.1 and 48.2 (ANOVA: *F*_2,106_ = 1886, *p* = <2e-16), post hoc *p*_48.1,48.2_ = <2e-04, *p*_48.1,48.4_ = <2e-16, *p*_48.2,48.4_ = <2e-16). In contrast, the high levels of variation within CCAMLR areas across the Southern Ocean preclude statistical significance across comparisons (ANOVA: *F*_2,21_ = 1.278, *p* = 0.299, post hoc *p*_48,58_ = 0.77, *p*_48,88_ = 0.42, *p*_58,88_ = 0.77), despite the observable trend towards reduced genome-wide diversity based on genotype-likelihoods in area 88 compared with areas 48 and 58 (Figure 5b), a finding corroborated by Watterson’s theta estimates of genomic diversity (Figure 5c). As with the admixture analysis, upon separation by cohort, a clearer pattern began to emerge, revealing a loss in heterozygosity over time (Figure 5d). Within management area 88, heterozygosity dropped significantly from the pre-2000 to the post-2000 cohort (ANOVA: *F*_1,11_ = 8.703, *p* = 0.0132, post hoc *p*_pre-2000,post-2000_ = 0.013), a loss also observed in Watterson’s theta estimates from area 88 (Figure 5e). This loss of diversity over time evidenced by genome-wide heterozygosity and Watterson’s theta estimates was also reflected in the 2× (*n* = 18, ANOVA: *F*_1,16_ = 11.9, *p* = 0.0033, post hoc *p*_pre-2000,post-2000_ = 0.0033) and noDS (*n* = 20, ANOVA: *F*_1,18_ = 7.909, *p* = 0.0115, post hoc *p*_pre-2000,post-2000_ = 0.012) datasets (Figure S7, Figure S9d, Figure S10d). Note that in management areas 48 and 58, post hoc *t*-tests between cohorts were not possible, because there was only one pre-2000 individual from these areas in all three datasets.

**Figure 5.**
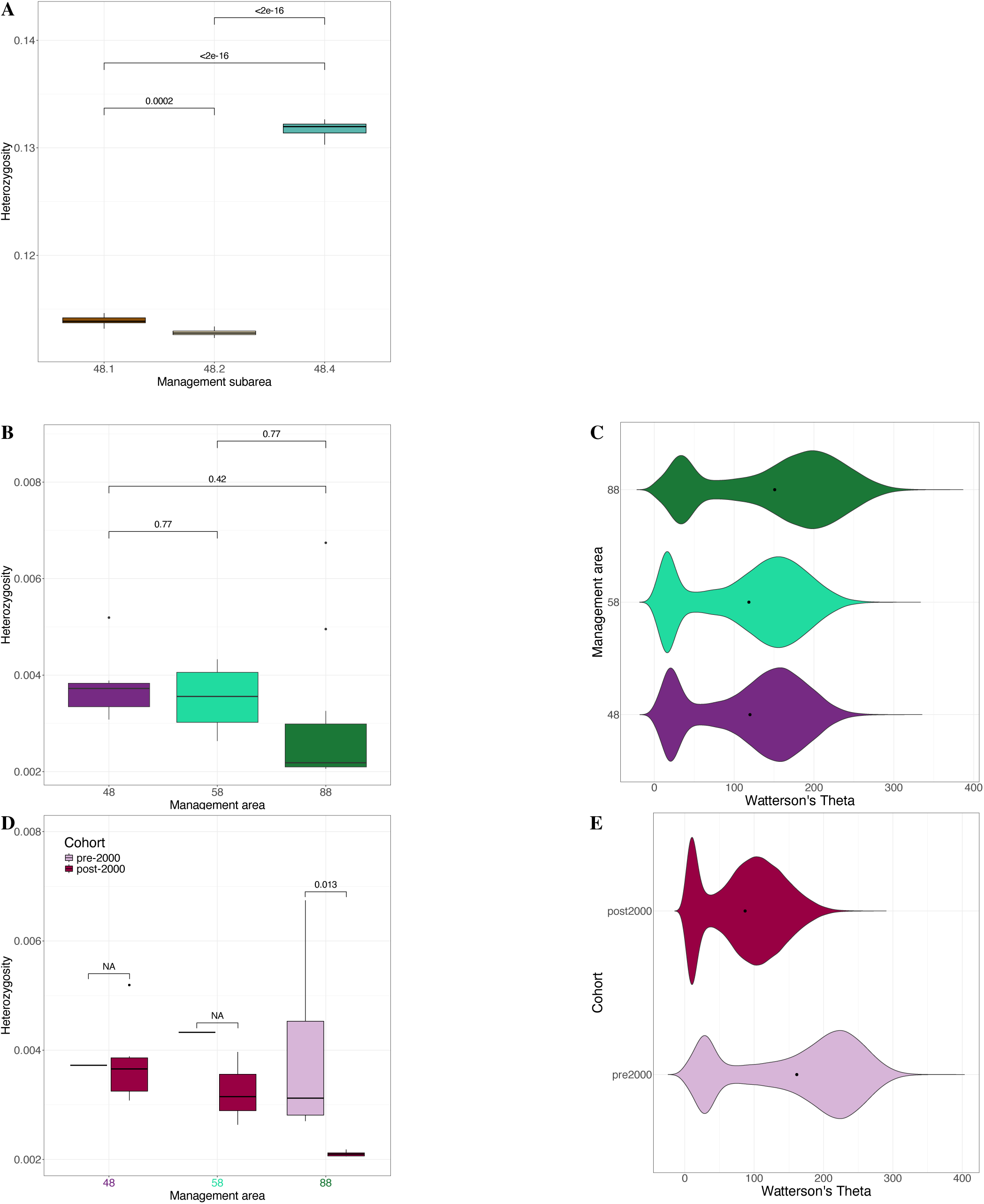
*D. mawsoni* genetic diversity based on (A) heterozygosity from 3RAD data (*n* = 108), and WGR data (*n* = 24) using (B) heterozygosity and (C) Watterson’s theta. Also shown: (D) heterozygosity and (E) Watterson’s theta between cohorts within area 88 (*n* = 13). Black dots = mean; NA = post hoc not possible.

Individual inbreeding coefficients (*F*) were significantly different between areas (Figure 7a), with area 88 exhibiting the highest *F* values (ANOVA: *F*_2,21_ = 7.003, *p* = 0.00468, post hoc *p*_48,58_ = 0.3012, *p*_48,88_ = 0.0038, *p*_58,88_ = 0.3012). To varying extents of significance, increased *F* values in area 88 persisted across the 2× (ANOVA: *F*_2,34_ = 8.788, *p* = 0.000839, post hoc *p*_48,58_ = 0.368, *p*_48,88_ = 0.001, *p*_58,88_ = 0.028) and noDS (ANOVA: *F*_2,38_ = 2.77, *p* = 0.0753, post hoc *p*_48,58_ = 1.00, *p*_48,88_ = 0.14, *p*_58,88_ = 0.17) datasets (Figure S8). Comparing *F* values between cohorts within area 88, the only area where such a comparison was possible, revealed no difference in *F* over time (ANOVA: *F*_1,11_ = 2.934, *p* = 0.115, post hoc *p*_pre-2000,post-2000_ = 0.11). This lack of change in *F* between cohorts was also observed in the 2× (ANOVA: *F*_1,16_ = 0.362, *p* = 0.556, post hoc *p*_pre-2000,post-2000_ = 0.56) and noDS (ANOVA: *F*_1,18_ = 0.142, *p* = 0.711, post hoc *p*_pre-2000,post-2000_ = 0.71) datasets. Runs of homozygosity (RoH) based on genotypes (and thus, only derived from the 5× dataset) supported evidence for increased inbreeding levels (F_ROH_) in CCAMLR management area 88 compared to areas 48 and 58 (Figure 7). F_ROH_ was significantly different between areas (Figure 7b), with once again area 88 showing the highest inbreeding levels (ANOVA: *F*_2,21_ = 6.144, *p* = 0.00793, post hoc *p*_48,58_ = 0.4747, *p*_48,88_ = 0.0091, *p*_58,88_ = 0.1296). As with *F*, there was no evidence of changes in F_ROH_ over time between cohorts (ANOVA: *F*_1,11_ = 0.146, *p* = 0.709, post hoc *p*_pre-2000,post-2000_ = 0.71).

Tajima’s D estimates derived from genotype-likelihoods were negative in all areas, though tended to be lower in CCAMLR management area 88 compared to areas 48 and 58 (Figure 6a), a trend reflected in the 2× (Figure S9a) and noDS (Figure S10a) datasets as well. While the influence of cohort was not possible to discern within areas 48 and 58 (given that there was only one pre-2000 individual from these areas in all three datasets), we were able to observe in area 88 a tendency towards higher Tajima’s D estimates in individuals from the post-2000 cohort compared to the pre-2000 cohort (Figure 6b), which persisted in the 2× (Figure S9c) and noDS (Figure S10c) datasets. Effective population size (*N_e_*) varied little over time, with *N_e_* in area 48(Figure 8) exhibiting a recent (<5000 years before present, yBP) drop after having remained nearly equivalent to that of area 58 previously. Area 88 consistently demonstrated the highest levels of *N_e_* among the three management areas.

**Figure 6.**
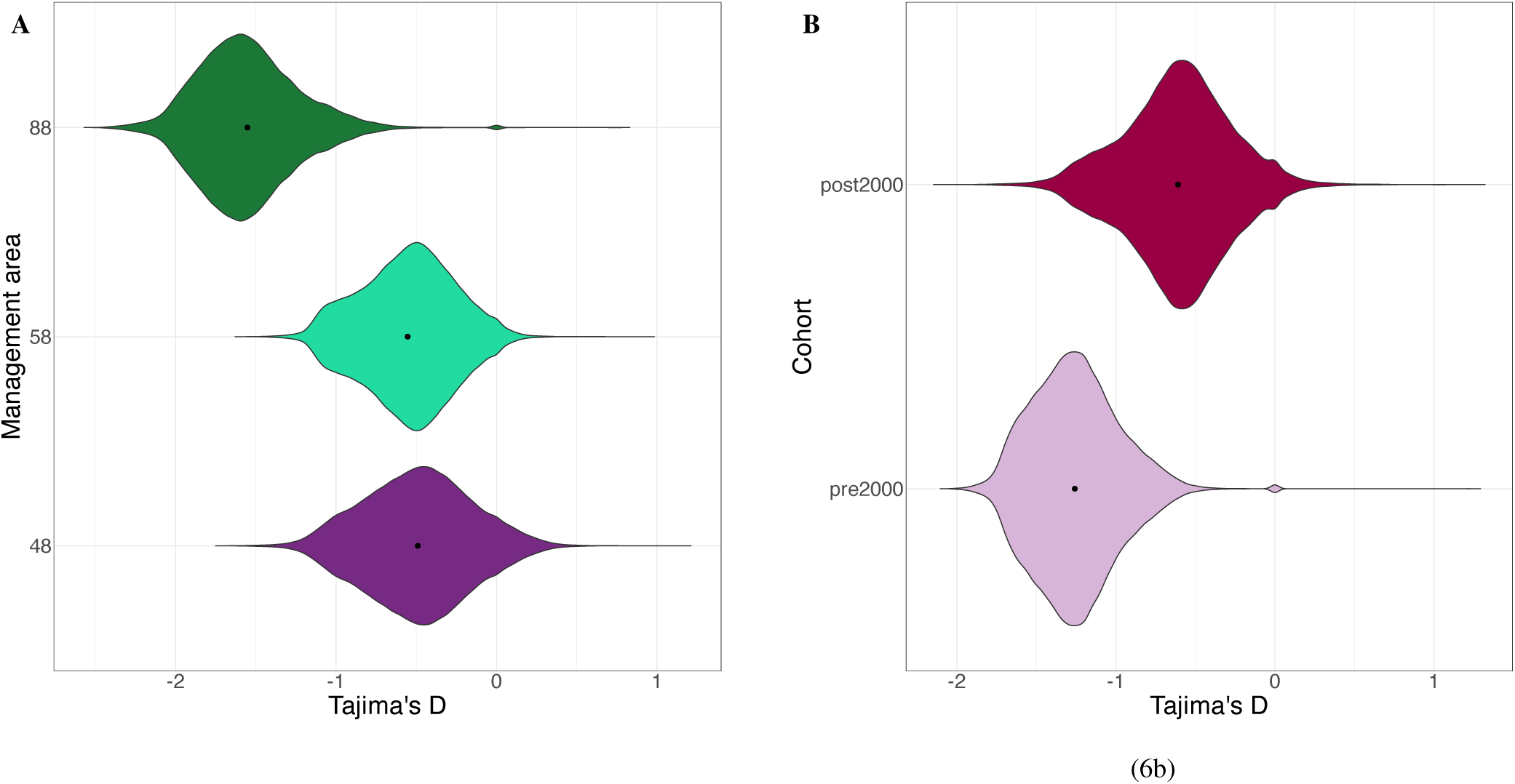
Violin plots of Tajima’s D derived from *D. mawsoni* WGR data using a sliding window approach with a window size of 50 kb and a step size of 10 kb; black dots indicate the mean value across the 50 kb-windows. (A) Tajima’s D across CCAMLR management areas (*n* = 24) and (B) between cohorts within area 88 (*n* = 13).

**Figure 7.**
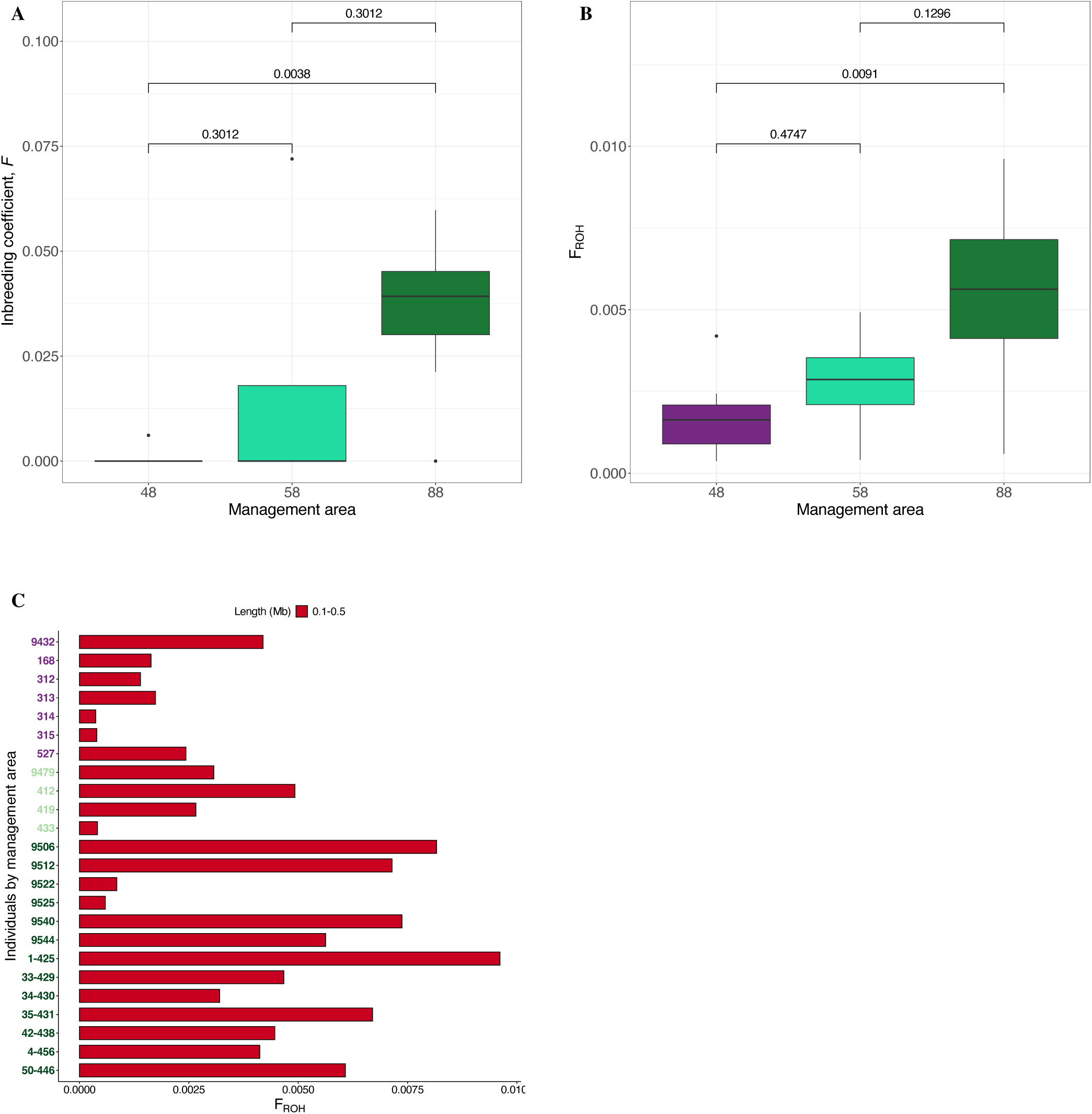
Inbreeding among *D. mawsoni* shown across CCAMLR management areas using (A) individual inbreeding coefficients, *F*, and (B & C) inbreeding level (FROH). *P*-values from post hoc pairwise *t*-tests are shown.

**Figure 8.**
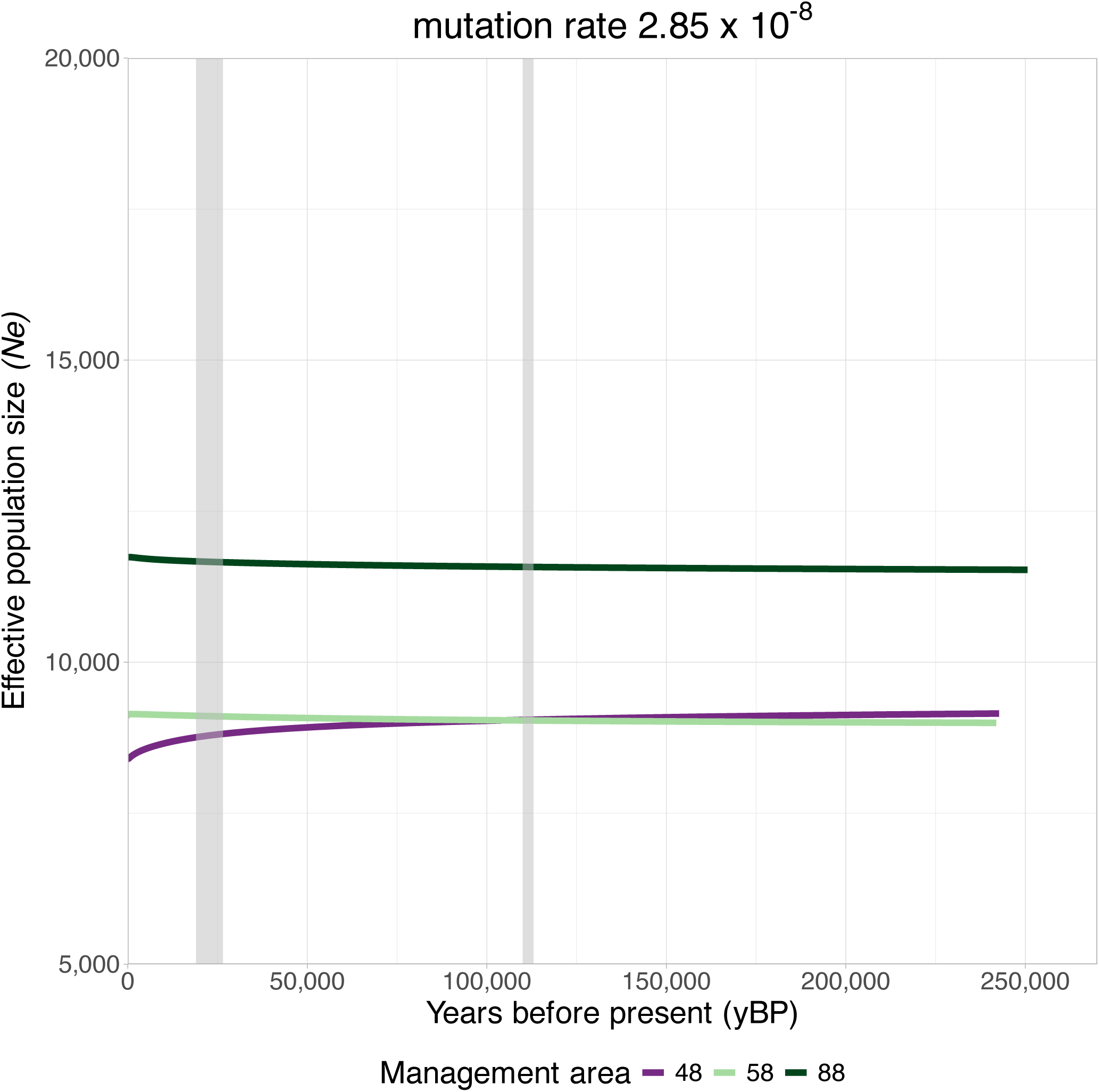
Effective population size (*Ne*) over time based on a generation time of 15 years and mutation rates of 2.85 × 10^−8^. The start of the last glacial period (110,000 yBP) and the last glacial maximum (26,500 - 19,000 yBP) are indicated by the gray bars.

## 4 Discussion

### 4.1 Population structure and demographic history of Antarctic toothfish

Our results generally support the panmixia hypothesis for *D. mawsoni* in the Southern Ocean, with some important exceptions. No population structure was evident from PCA and admixture analyses both based on 3RADseq libraries from the Weddell Sea (CCAMLR management subareas 48.1, 48.2 and 48.4) (Figure 3) as well as WGR libraries from circumpolar distributions (CCAMLR management areas 48, 58 and 88) (Figure 4). This is consistent with the results of PCA and Principle Coordinate Analysis in Ceballos et al. (2021) and Maschette et al. (2023) respectively, as well as the factorial correspondence analysis in Choi et al. (2021).

While admixture results across management areas further supported a lack of geographic genetic structure (Figure 4b), when considered across cohort, the pre-2000 cohort showed evidence of reduced admixture, while those from the post-2000 cohort were nearly all admixed (Figure 4c). The increased levels of admixture observed in the post-2000 cohort may be related to the influx of new genotypes from one or more source populations (Fitzpatrick et al., 2020) following reductions in IUU fishing for toothfish after the 1990s (Agnew, 2000; Miller, 2004; Kock, 2007; Baird, 2024). We also observed a trend towards reduced heterozygosity in area 88 (Figure 5b), which was revealed to be driven by a significant drop in heterozygosity between the pre-2000 and post-2000 cohorts in that area (Figure 5d). The loss of genetic diversity over time in area 88 was further supported by the discovery of elevated inbreeding evidenced both by the significant increase in *F* (Figure 7a) and F_ROH_ (Figure 7), as well as the drop in Watterson’s Theta (Figure 5e). This loss in heterozygosity would seem to contradict the increase in admixture that we observe. However, it is possible that if the source populations contributing to the influx of new genotypes promoting admixture in area 88 had lower heterozygosity than the resident population, this could drive a decrease in heterozygosity concurrent with increased admixture (Boca et al., 2020). Furthermore, there is evidence from human populations of increased admixture coincident with inbreeding (Yang et al., 2014). Indeed, low recruitment years (i.e., periods when proportionally few mature adults reproduce), may spur increases in inbreeding (O’Leary et al., 2013). While Tajima’s D is reduced in area 88 with respect to areas 48 and 58 (Figure 6a), this may ultimately be driven by reduced Tajima’s D in the pre-2000 cohort (Figure 6b). Increased levels of Tajima’s D in the post-2000 cohort in area 88 with respect to the pre-2000 cohort may be evident of a population contraction between the cohorts (Figure 6b), which is supported by the reductions in heterozygosity over time in area 88 (Figure 5d).

The lack of change in *N*_e_ over time for areas 58 and 88 (Figure 8) differs from the slight recent drop observed in area 48 (Figure 8). These observations could have two explanations: 1) SMC++ has a tendency to smooth over abrupt changes in *N*_e_ over time, which could cause the observed lack of change (Terhorst et al., 2017); and 2) the recent declines observed in *N*_e_ could be caused either by population substructure, or by population expansions (Bansal and Nichols, 2025). It is thus difficult to draw conclusions regarding *N*_e_ in *D. mawsoni* over this time scale. Indeed, estimating *N*_e_ has historically been a challenge in large marine populations (Waples, 2016); future work using larger sample sizes, and multiple approaches to estimating *N*_e_ should help to resolve the question of demographic changes *D. mawsoni* across time scales (Delord et al., 2025).

### 4.2 Drivers of Antarctic toothfish population dynamics

The two key exceptions to panmixia identified in *D. mawsoni* are thus: 1) over time, genetic diversity is reduced, admixture increases, and populations contract; and 2) in CCAMLR management area 88, genetic diversity has reduced while populations have recently expanded. Among the possible explanations for these observations, two possibilities stand out: 1) chaotic genetic patchiness; and 2) fisheries pressure. Chaotic genetic patchiness, or changes in genetic diversity over time linked to recruitment variability (Johnson and Black, 1982; Eldon et al., 2016), has been observed in Antarctic fish (Papetti et al., 2007, 2012), and in particular in a key prey species of *D. mawsoni*, *Pleuragramma antarctica* (Agostini et al., 2015; Caccavo et al., 2018). Indeed, recruitment success in *D. mawsoni* has been linked to climate variability driven by the impact on sea ice drift of the Amundsen Sea Low, whose position and strength is modulated by the El Niño-Southern Oscillation (ENSO) (Behrens et al., 2021, 2024). Indeed, Behrens et al. (2024) identify recruitment anomalies (i.e., changes in the number of individuals available to the fishery in a given year) in *D. mawsoni* between 1995 - 2012 in area 88, which may explain the reduced diversity and population contractions observed.

Because fishing pressure can also result in a loss of genetic diversity (Pinsky and Palumbi, 2014; Sadler et al., 2023; Aquije et al., 2024), IUU fishing for *D. mawsoni* during the 1990s (Österblom and Sumaila, 2011; Lack, 2008; Ainley et al., 2012; Brooks, 2013) could also explain the loss of genetic diversity observed in the post-2000 cohort. However, the greatest levels of IUU fishing occurred outside of area 88 (Österblom and Sumaila, 2011), which would not account for the greater reductions in genetic diversity observed in this area. CCAMLR management area 88 has historically experienced the highest levels of legal fishing since the start of the fishery in the late 1990s (Figure 2b) (Kock et al., 2007). It is thus possible that management practices in area 88, while meeting the CCAMLR’s standards for sustainability based on regular stock assessments (Hanchet et al., 2015b), are ultimately not precautionary, and are driving genetic erosion (Leroy et al., 2018) in *D. mawsoni* populations of area 88.

### 4.3 Implications for fisheries management

Both recruitment variability and fisheries pressure likely play a synergistic role in modulating gene flow within and between management areas of *D. mawsoni* (Caccavo et al., 2021). To tease apart the role of these two factors, future research must: 1) regularly monitor genetic diversity in *D. mawsoni* using standardized approaches to allow for consistency between years; and 2) increase sample sizes across cohorts in all regions to better understand the relationship between genetic diversity and recruitment anomalies on a circumpolar scale. The method established by our research group to create WGR libraries from DNA extracted from otoliths (Caccavo et al., 2024) holds promise as a cheap, low time-investment, and widely applicable approach to monitoring *D. mawsoni* genetic diversity beyond assessments of gene flow between management areas, which do not capture demographic changes critical to ensuring the sustainability of the fishery. Many examples now exist of fisheries that have not only aspired to integrate genetics in their management strategies, but have succeeded in doing so (Derycke and Maes, 2023), including the example of the Atlantic herring (*Clupea harengus*), in which whole-genome resequencing illuminated previously undetectable genetic structure and population dynamics across north Atlantic stocks (Lamichhaney et al., 2017; Han et al., 2020).

Ultimately, fisheries management of *D. mawsoni* will have to move beyond proxies of diversity and connectivity to integrate genetic resilience into stock assessment models. Given their early-life dependence on sea ice (Parker et al., 2021), which has experienced historic perturbations in the Southern Ocean in recent years (Josey et al., 2024; Eayrs et al., 2019; Liu et al., 2023; Turner et al., 2022), it will be critical for future research to understand the environmental variables driving climate change impacts on *D. mawsoni*. Indeed, the >121 thousand putatively adaptive loci that we identified among genotypes derived from the 5× dataset could be used to apply climate genomics (Caccavo et al., 2023) approaches to *D. mawsoni*. Indeed, from a fisheries management perspective, adaptation signals a response to environmental change that can leave a population vulnerable to further pressures from exploitation, given that particularity in fish species, adaptive evolution almost never results in the fixation of beneficial alleles (Bernatchez, 2016). Ultimately, climate genomics approaches that include projections of future environmental change, including future gSDM (Fiscus et al., 2025) and genomic offset (Fitzpatrick and Keller, 2015; Layton et al., 2024), will allow fisheries managers to integrate assessments of long-term genomic change in *D. mawsoni* into more precautionary and fundamentally sustainable management strategies.

## Data availability

Raw sequencing reads are available from GenBank under BioProject number PRJNA1366982 for 3RADseq reads, and BioProject number PRJNA1123220 for WGR reads, with associated samples and sequences deposited in BioSample, and Sequence Read Archive. Data files, commands, and scripts for various bioinformatics analyses are accessible from a dedicated GitHub repository.

## Author contributions

JAC, LSA and CJM conceived of and designed the research. JAC facilitated the acquisition of samples. JAC performed the laboratory work and conducted the bioinformatics analyses with support from LSA, SM and SP. EC provided support for the preprocessing of the high-output data, SNP calling, and various downstream analyses. JAC drafted the manuscript with input from all authors. All authors read and approved the manuscript.

## Acknowledgements

We thank Steve Parker (New Zealand), Illia Slypko (Ukraine), Phil Hollyman (United Kingdom), and Patrick Gaffney (United States) for providing tissue samples used to build the 3RADseq libraries. This work was completed while JAC was receiving support from the Alexander von Humboldt Foundation in the form of a Humboldt Research Fellowship for Postdoctoral Researchers. JAC now acknowledges support from a postdoctoral fellowship adminstered by the Institut Pierre-Simon Laplace (IPSL), made possible by French state aid managed by the ANR (‘Agence Nationale de la Recherche’) under the ‘Investissements d’avenir’ program, with the reference ANR-11-IDEX-0004-17-EURE-0006.

## Supplementary Tables

**Table S1.** Group sizes (*n*) across datasets.

| 3RAD dataset | subarea 48.1 | subarea 48.2 | subarea 48.4 |
| --- | --- | --- | --- |
| Subarea totals | 40 | 41 | 28 |
| Dataset total |  |  | 108 |
| 5× WGR dataset | area 48 | area 58 | area 88 |
| pre-2000 | 1 | 1 | 6 |
| post-2000 | 6 | 3 | 7 |
| Area totals | 7 | 4 | 13 |
| Dataset total |  |  | 24 |
| 2× WGR dataset | area 48 | area 58 | area 88 |
| pre-2000 | 1 | 1 | 10 |
| post-2000 | 10 | 7 | 8 |
| Area totals | 11 | 8 | 18 |
| Dataset total |  |  | 37 |
| noDS WGR dataset | area 48 | area 58 | area 88 |
| pre-2000 | 1 | 1 | 11 |
| post-2000 | 12 | 7 | 9 |
| Area totals | 13 | 8 | 20 |
| Dataset total |  |  | 41 |

**Table S2.** Sample metadata including type, management area and subarea, latitude, longitude, year of capture, estimated year of birth, cohort, total fish length (cm), final coverage, and dataset inclusion. 2×, downsampled to 2×; 5×, downsampled to 5×; noDS, not downsampled; na, not available.

| Sample | Type | Area | Subarea | Latitude | Longitude | Year of capture | Estimated birth year | Cohort | Length (cm) | Coverage | Dataset inclusion |
| --- | --- | --- | --- | --- | --- | --- | --- | --- | --- | --- | --- |
| 9432 | tissue | 48 | 48.1 | -62.00 | -58.00 | 2001 | 1985 | pre-2000 | na | 11.40 | noDS, 2×, 5× |
| 1 | otolith | 48 | 48.1 | -63.24 | -59.57 | 2006 | 2000 | pre-2000 | 54.0 | 3.40 | noDS, 2× |
| 312 | otolith | 48 | 48.5 | -74.64 | -27.84 | 2013 | 2000 | post-2000 | 123.0 | 12.46 | noDS, 2×, 5× |
| 313 | otolith | 48 | 48.5 | -74.64 | -27.84 | 2013 | 2000 | post-2000 | 118.0 | 5.92 | noDS, 2×, 5× |
| 314 | otolith | 48 | 48.5 | -74.64 | -27.84 | 2013 | 2000 | post-2000 | 103.0 | 13.27 | noDS, 2×, 5× |
| 315 | otolith | 48 | 48.5 | -74.64 | -27.84 | 2013 | 2000 | post-2000 | 128.0 | 10.86 | noDS, 2×, 5× |
| 527 | otolith | 48 | 48.2 | -61.56 | -38.72 | 2018 | 2005 | post-2000 | 151.0 | 6.76 | noDS, 2×, 5× |
| 67 | otolith | 48 | 48.1 | -62.08 | -53.55 | 2019 | 2005 | post-2000 | 141.0 | 4.23 | noDS, 2× |
| 95 | otolith | 48 | 48.1 | -62.16 | -53.57 | 2019 | 2005 | post-2000 | 140.0 | 1.13 | noDS |
| 164 | otolith | 48 | 48.2 | -61.40 | -34.74 | 2019 | 2005 | post-2000 | 156.0 | 2.94 | noDS, 2× |
| 168 | otolith | 48 | 48.2 | -61.40 | -34.74 | 2019 | 2005 | post-2000 | 158.0 | 6.92 | noDS, 2×, 5× |
| 231 | otolith | 48 | 48.4 | -58.58 | -26.12 | 2019 | 2005 | post-2000 | 116.0 | 1.81 | noDS |
| 251 | otolith | 48 | 48.4 | -58.77 | -25.25 | 2019 | 2005 | post-2000 | 167.0 | 3.04 | noDS, 2× |
| 9479 | tissue | 58 | 58.4.2 | -67.20 | 44.00 | 2001 | 1985 | pre-2000 | na | 6.43 | noDS, 2×, 5× |
| 398 | otolith | 58 | 58.4.2 | -66.56 | 71.95 | 2019 | 2010 | post-2000 | 86.0 | 3.59 | noDS, 2× |
| 402 | otolith | 58 | 58.4.2 | -66.56 | 71.95 | 2019 | 2010 | post-2000 | 70.0 | 4.84 | noDS, 2× |
| 408 | otolith | 58 | 58.4.2 | -66.58 | 71.68 | 2019 | 2005 | post-2000 | 136.5 | 2.46 | noDS, 2× |
| 412 | otolith | 58 | 58.4.2 | -66.49 | 71.74 | 2019 | 2010 | post-2000 | 105.0 | 6.88 | noDS, 2×, 5× |
| 419 | otolith | 58 | 58.4.2 | -66.60 | 71.48 | 2019 | 2010 | post-2000 | 89.5 | 12.03 | noDS, 2×, 5× |
| 433 | otolith | 58 | 58.4.2 | -66.44 | 71.60 | 2019 | 2005 | post-2000 | 122.5 | 11.07 | noDS, 2×, 5× |
| 435 | otolith | 58 | 58.4.2 | -66.44 | 71.60 | 2019 | 2010 | post-2000 | 98.0 | 3.28 | noDS, 2× |
| 9506 | tissue | 88 | 88.1 | -77.00 | 179.00 | 2001 | 1985 | pre-2000 | na | 7.29 | noDS, 2×, 5× |
| 9512 | tissue | 88 | 88.1 | -77.00 | 179.00 | 2001 | 1985 | pre-2000 | na | 22.13 | noDS, 2×, 5× |
| 9522 | tissue | 88 | 88.1 | -73.00 | 171.00 | 2001 | 1985 | pre-2000 | na | 9.52 | noDS, 2×, 5× |
| 9525 | tissue | 88 | 88.1 | -71.00 | 176.00 | 2001 | 1985 | pre-2000 | na | 7.94 | noDS, 2×, 5× |

Table S2 (continued)
| Sample | Type | Area | Subarea | Latitude | Longitude | Year of capture | Estimated birth year | Cohort | Length (cm) | Coverage | Dataset inclusion |
| --- | --- | --- | --- | --- | --- | --- | --- | --- | --- | --- | --- |
| 9540 | tissue | 88 | 88.1 | -70.00 | 179.00 | 2001 | 1985 | pre-2000 | na | 13.40 | noDS, 2×, 5× |
| 9544 | tissue | 88 | 88.1 | -70.00 | 179.00 | 2001 | 1985 | pre-2000 | na | 5.90 | noDS, 2×, 5× |
| 12821 | tissue | 88 | 88.2 | -68.00 | -122.00 | 2005 | 1990 | pre-2000 | na | 3.43 | noDS, 2× |
| 12822 | tissue | 88 | 88.2 | -68.00 | -122.00 | 2005 | 1990 | pre-2000 | na | 1.76 | noDS |
| 12836 | tissue | 88 | 88.2 | -68.00 | -122.00 | 2005 | 1990 | pre-2000 | na | 2.32 | noDS, 2× |
| 12840 | tissue | 88 | 88.2 | -68.00 | -123.00 | 2005 | 1990 | pre-2000 | na | 2.29 | noDS, 2× |
| 12841 | tissue | 88 | 88.2 | -68.00 | -123.00 | 2005 | 1990 | pre-2000 | na | 2.77 | noDS, 2× |
| 1-425 | tissue | 88 | 88.1 | -64.00 | 178.00 | 2019 | 2005 | post-2000 | 148.0 | 15.37 | noDS, 2×, 5× |
| 33-429 | tissue | 88 | 88.1 | -72.00 | 181.00 | 2019 | 2005 | post-2000 | 156.0 | 13.09 | noDS, 2×, 5× |
| 34-430 | tissue | 88 | 88.1 | -72.00 | 181.00 | 2019 | 2005 | post-2000 | 142.0 | 11.82 | noDS, 2×, 5× |
| 35-431 | tissue | 88 | 88.1 | -72.00 | 181.00 | 2019 | 2005 | post-2000 | 152.0 | 8.78 | noDS, 2×, 5× |
| 4-456 | tissue | 88 | 88.1 | -65.00 | 176.00 | 2019 | 2005 | post-2000 | 145.0 | 19.92 | noDS, 2×, 5× |
| 6-450 | tissue | 88 | 88.1 | -65.00 | 178.00 | 2019 | 2005 | post-2000 | 142.0 | 2.88 | noDS, 2× |
| 38-434 | tissue | 88 | 88.1 | -73.00 | 180.00 | 2020 | 2005 | post-2000 | 134.0 | 1.72 | noDS |
| 42-438 | tissue | 88 | 88.1 | -73.00 | 180.00 | 2020 | 2005 | post-2000 | 127.0 | 11.28 | noDS, 2×, 5× |
| 50-446 | tissue | 88 | 88.1 | -73.00 | 180.00 | 2020 | 2005 | post-2000 | 158.0 | 27.09 | noDS, 2×, 5× |

## Supplementary Figures

**Figure S1.**
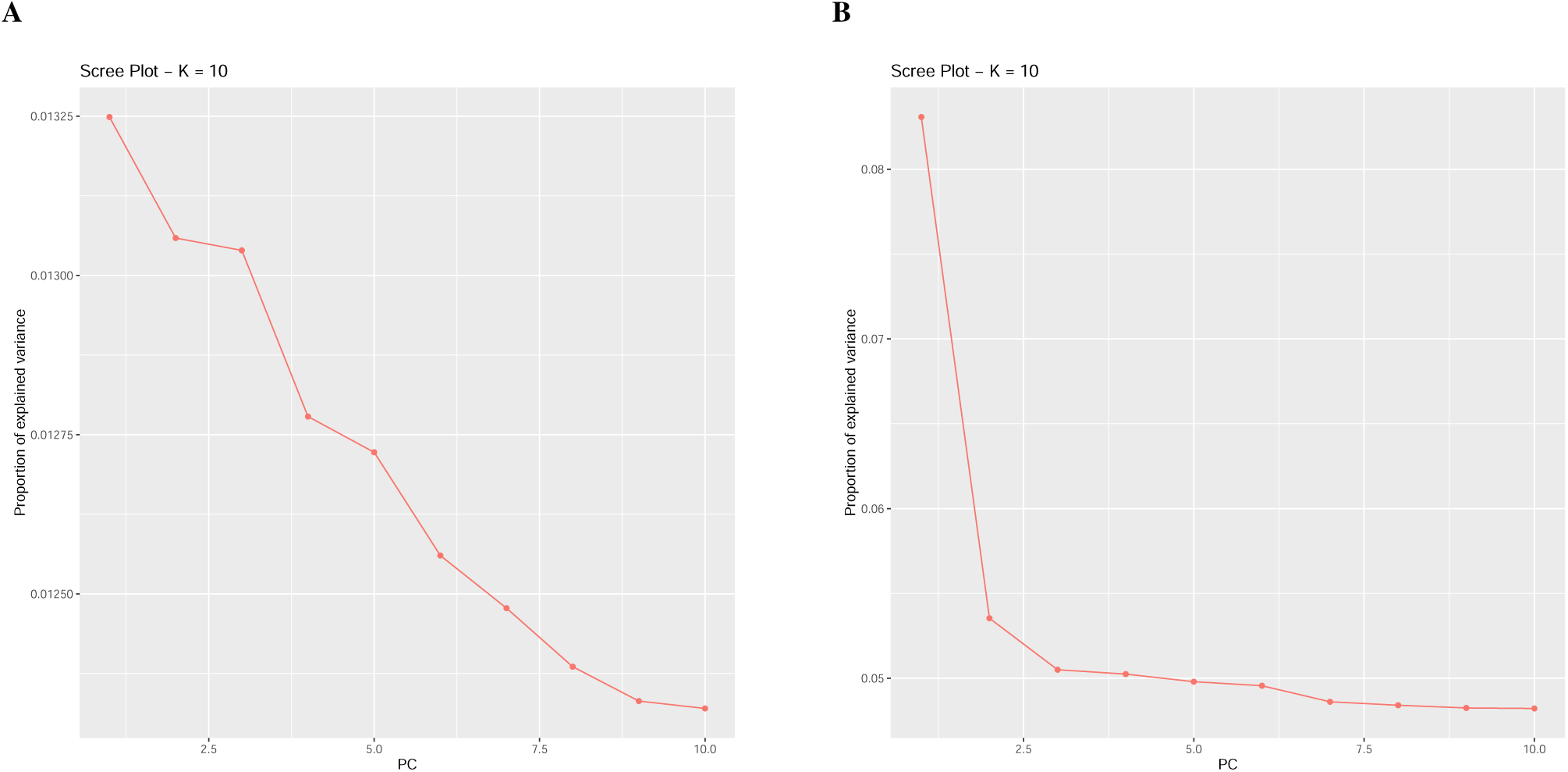
Scree plots generated in PCADAPT supporting the use of 2 principal components (K) for outlier scans based on (A) 3RADseq and (B) WGR data (adapted from Caccavo et al. (2024), Figure S1).

**Figure S2.**
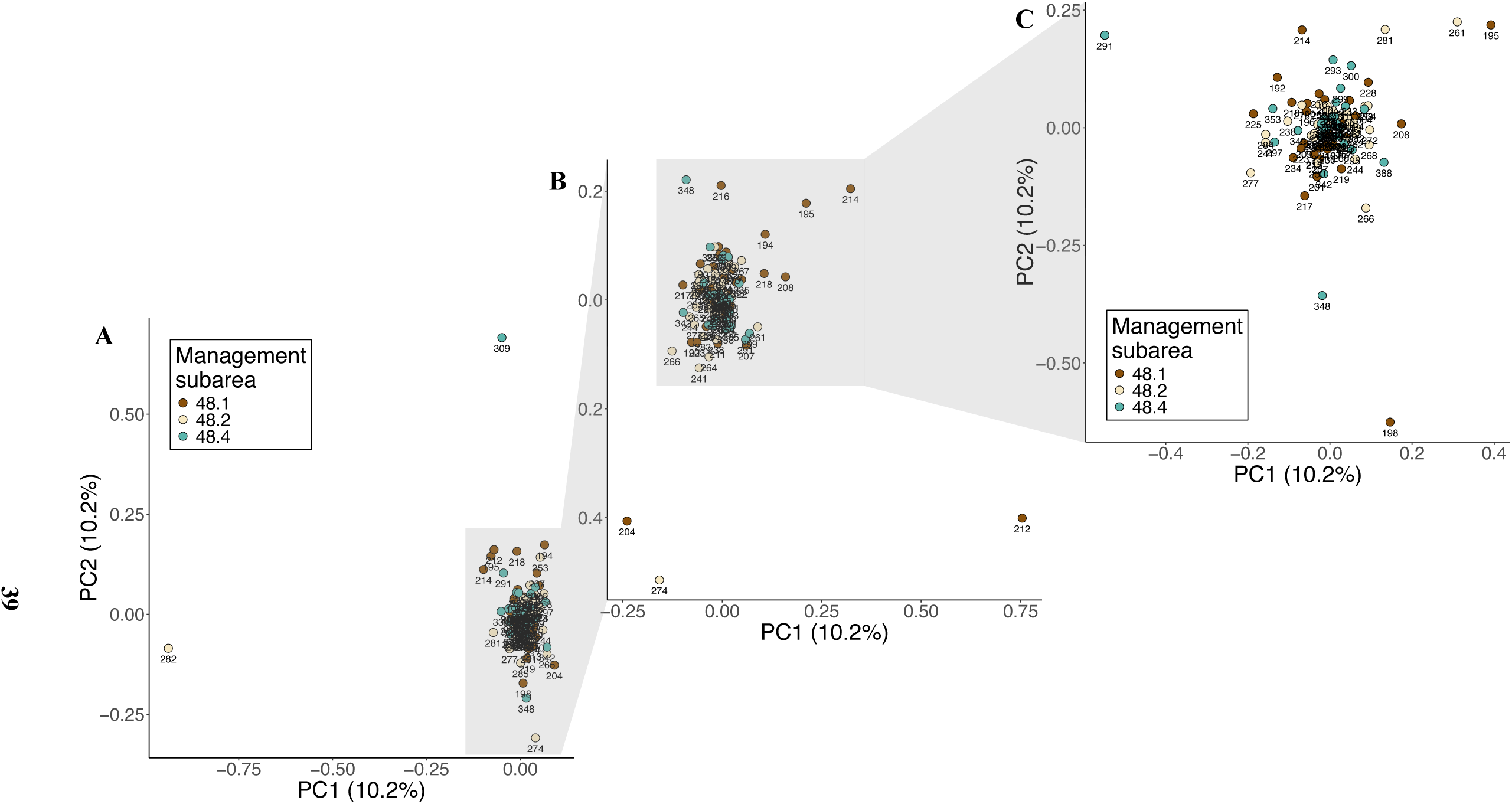
PCA based on 3RADseq data of *D. mawsoni* in the Weddell Sea. Groups labeled by CCAMLR management subarea (48.1, 48.2, 48.3). (A) Original PCA *n* = 109 samples seen in Figure 3a highlighting samples under the gray box used in the subsequent PCA of (B) *n* = 106 samples, with 2 outliers (samples 282 and 309) removed, highlighting samples under the gray box used in the final PCA of (C) *n* = 103 samples, with an additional 3 outliers removed (samples 282 and 309 as well as samples 204, 274, and 212). Data labels indicate sample IDs.

**Figure S3.**
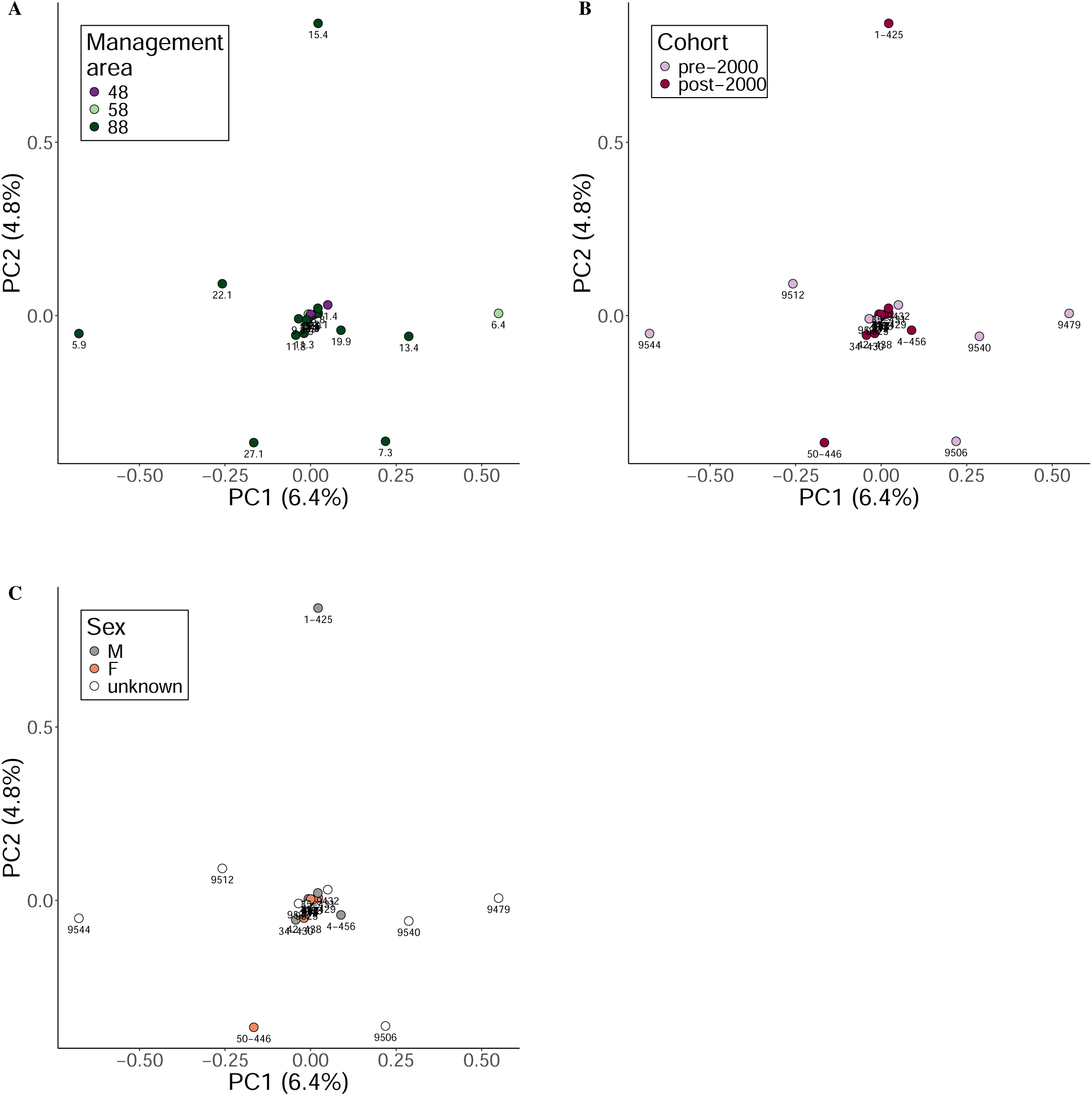
PCA plots of alternative sample groupings based on WGR data. (A) CCAMLR management area (48, 58, 88), data labels indicate coverage (×). (B) Cohort, data labels indicate sample ID. (C) Sex, data labels indicate sample ID.

**Figure S4.**
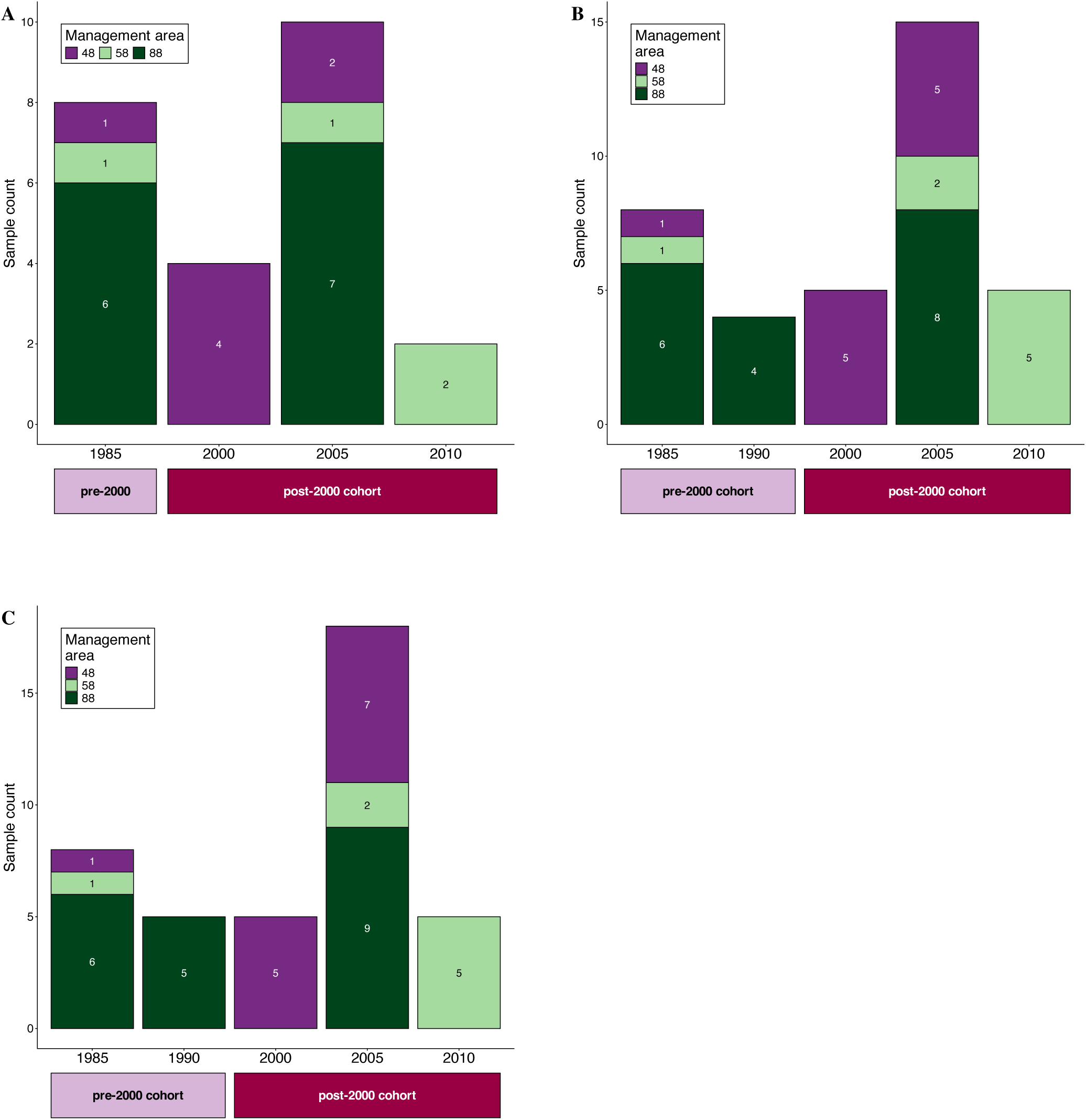
Sample counts across cohorts and management areas from (A) 5× (*n* = 24), (B) 2× (*n* = 37) and (C) noDS (*n* = 41) datasets.

**Figure S5.**
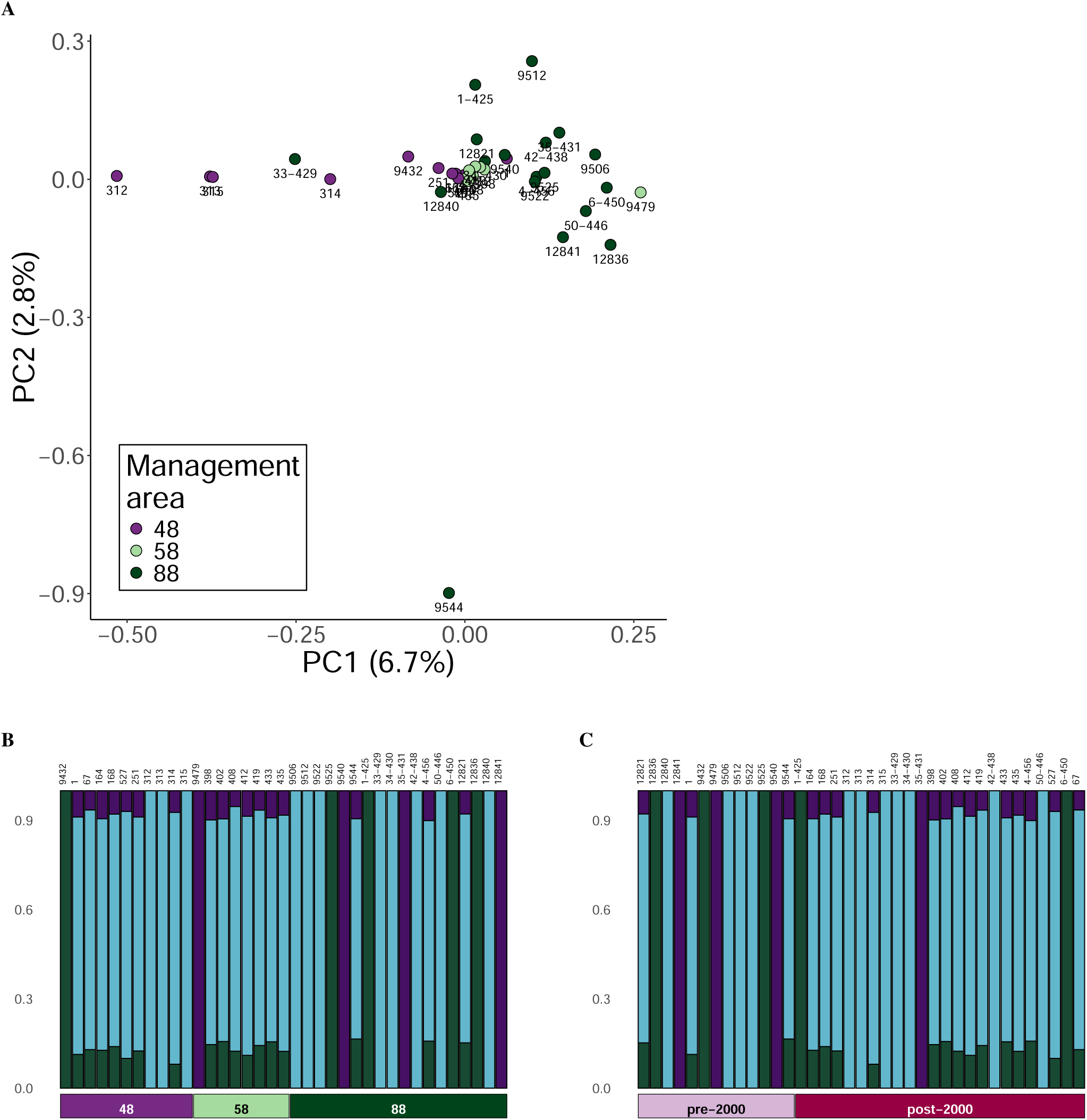
Circumpolar population structure of *D. mawsoni* based on WGR data (2× dataset, *n* = 37). Groups labeled by CCAMLR management areas (48, 58, 88). (A) PCA based on all samples, data labels indicate sample IDs. Admixture plot of population structure for *K* = 3, with sample IDs listed above the plot; ordered by (B) CCAMLR area. (C) cohort.

**Figure S6.**
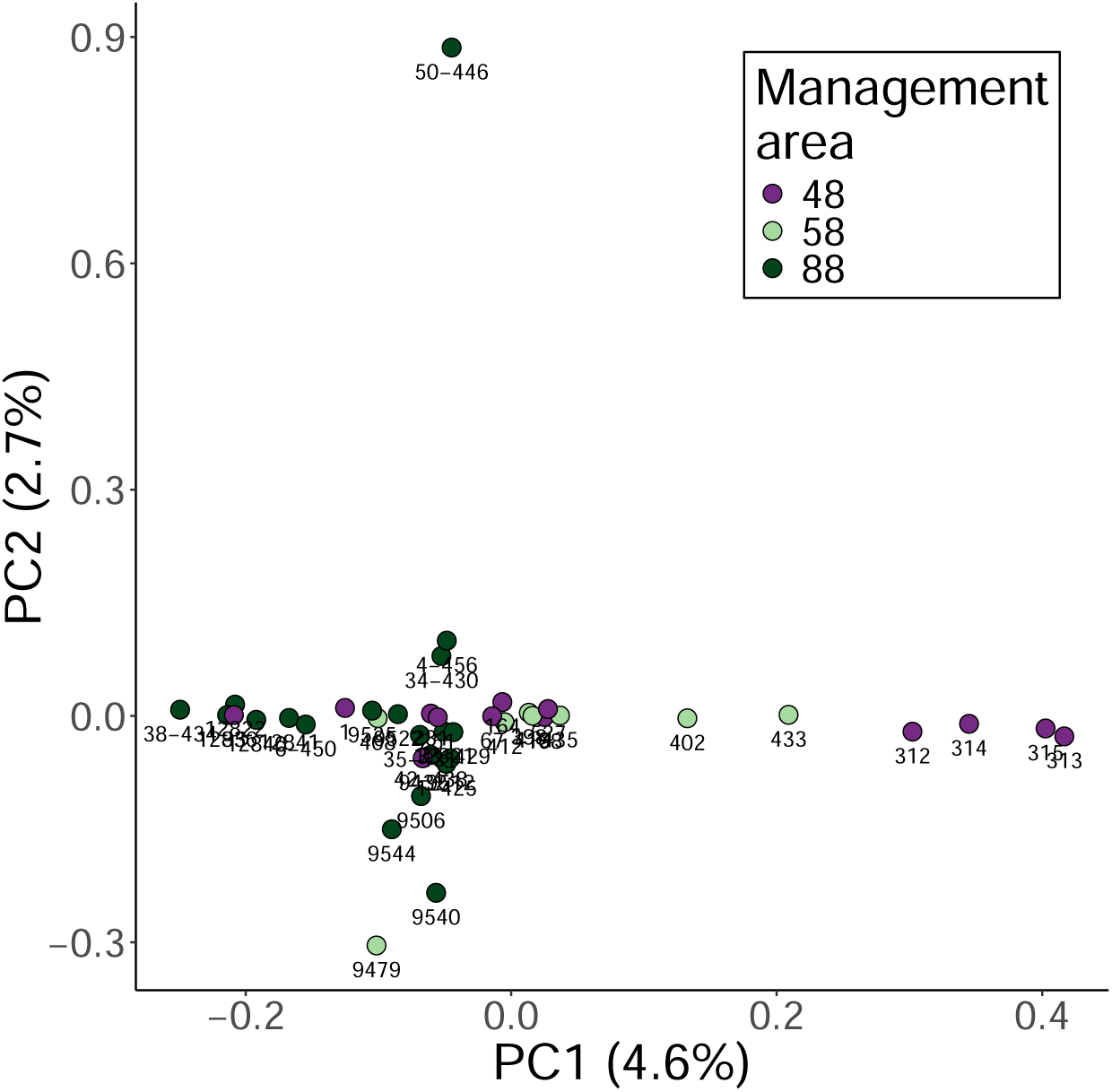
Circumpolar population structure of *D. mawsoni* based on WGR data (noDS dataset, *n* = 41). Groups labeled by CCAMLR management areas (48, 58, 88). PCA based on all samples, data labels indicate sample IDs.

**Figure S7.**
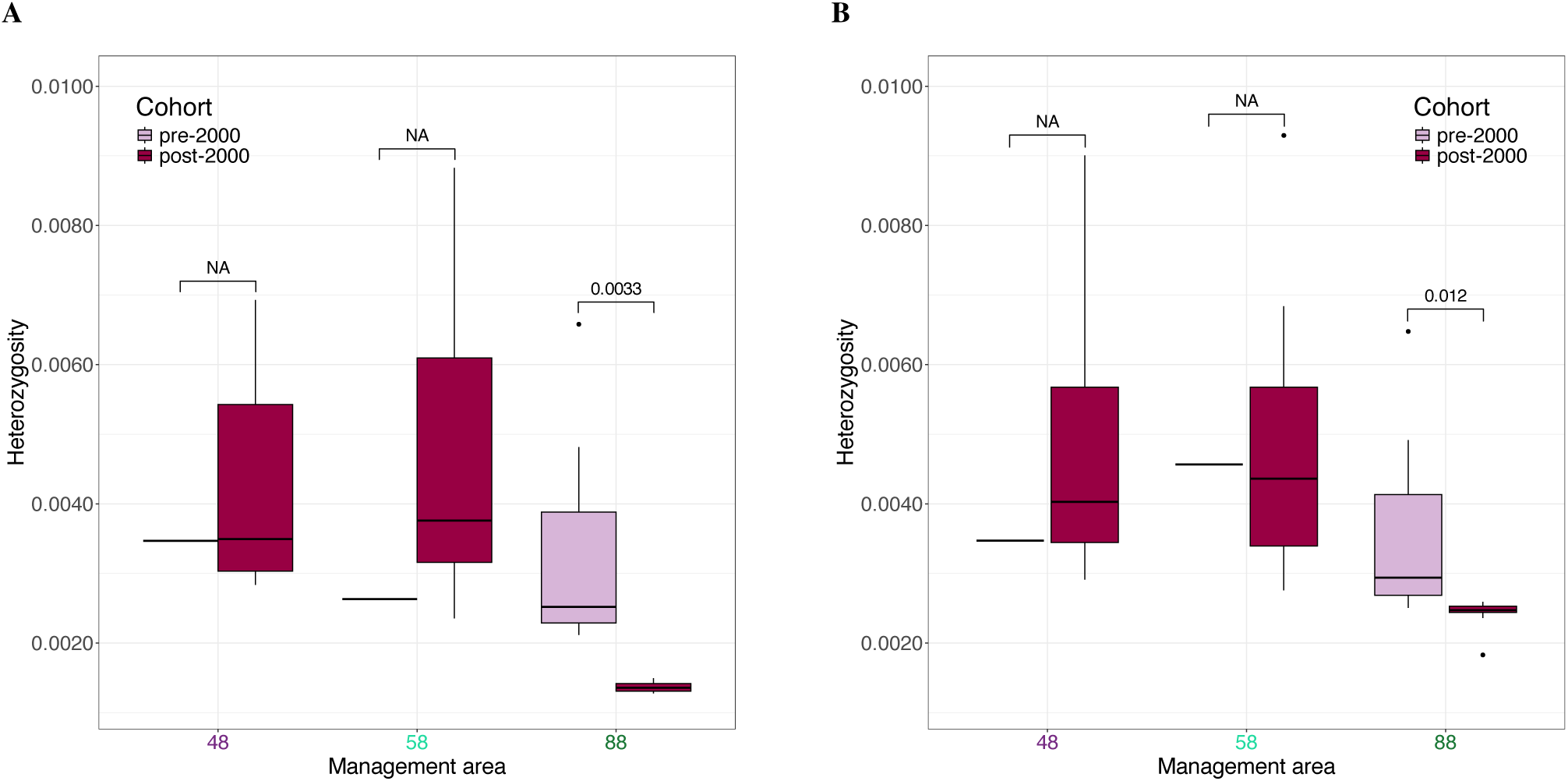
*D. mawsoni* heterozygosity from all three circumpolar management areas including across cohorts based on (A) 2× (*n* = 37) and (B) noDS (*n* = 41) datasets. *P*-values from post hoc pairwise *t*-tests are shown. Where NA indicated, post hoc tests were not possible due to groups with *n* = 1 individual.

**Figure S8.**
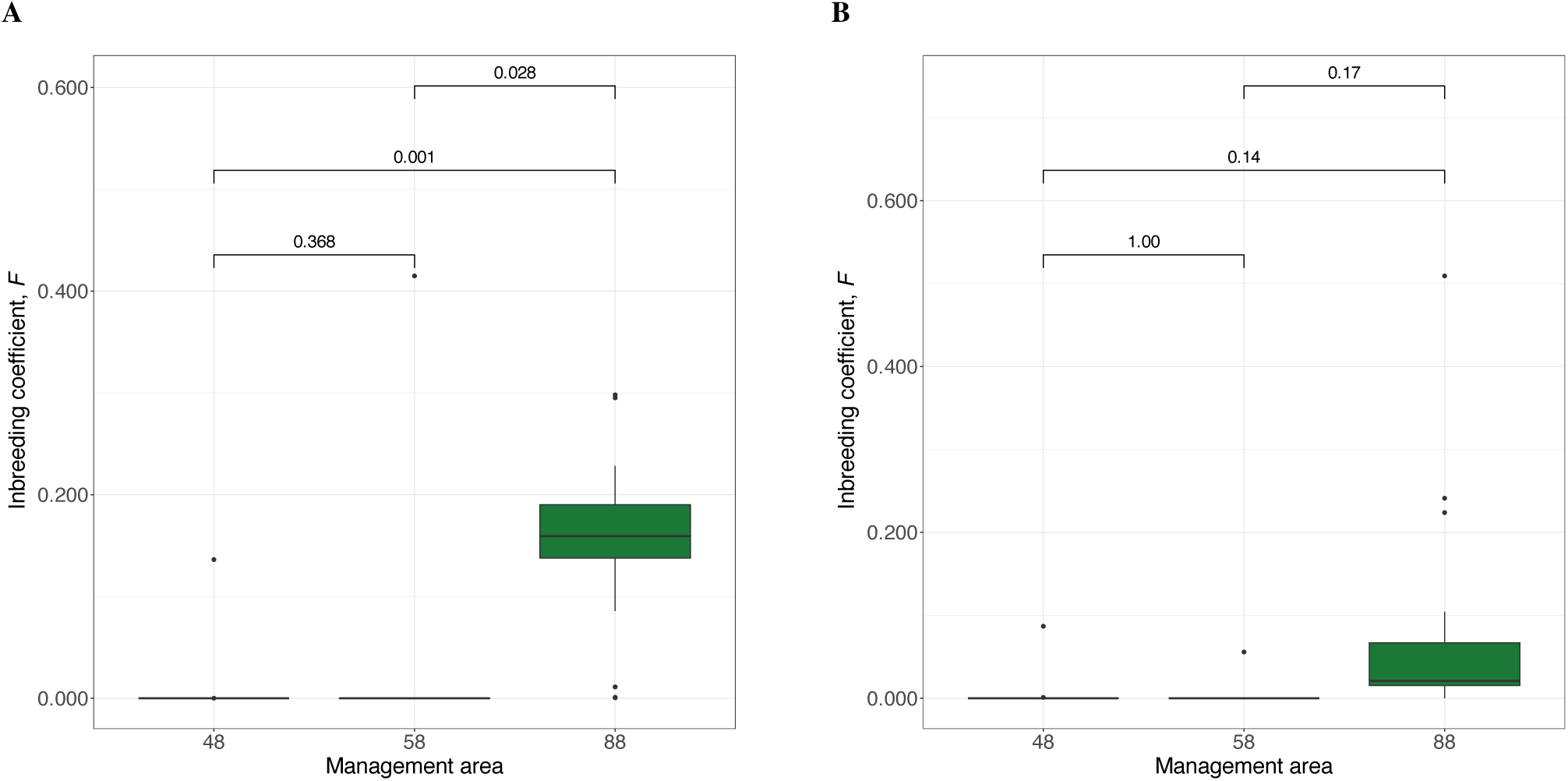
*D. mawsoni* individual inbreeding coefficients, *F*, based on (A) 2× (*n* = 37) and (B) noDS (*n* = 41) datasets. *P*-values from post hoc pairwise *t*-tests are shown.

**Figure S9.**
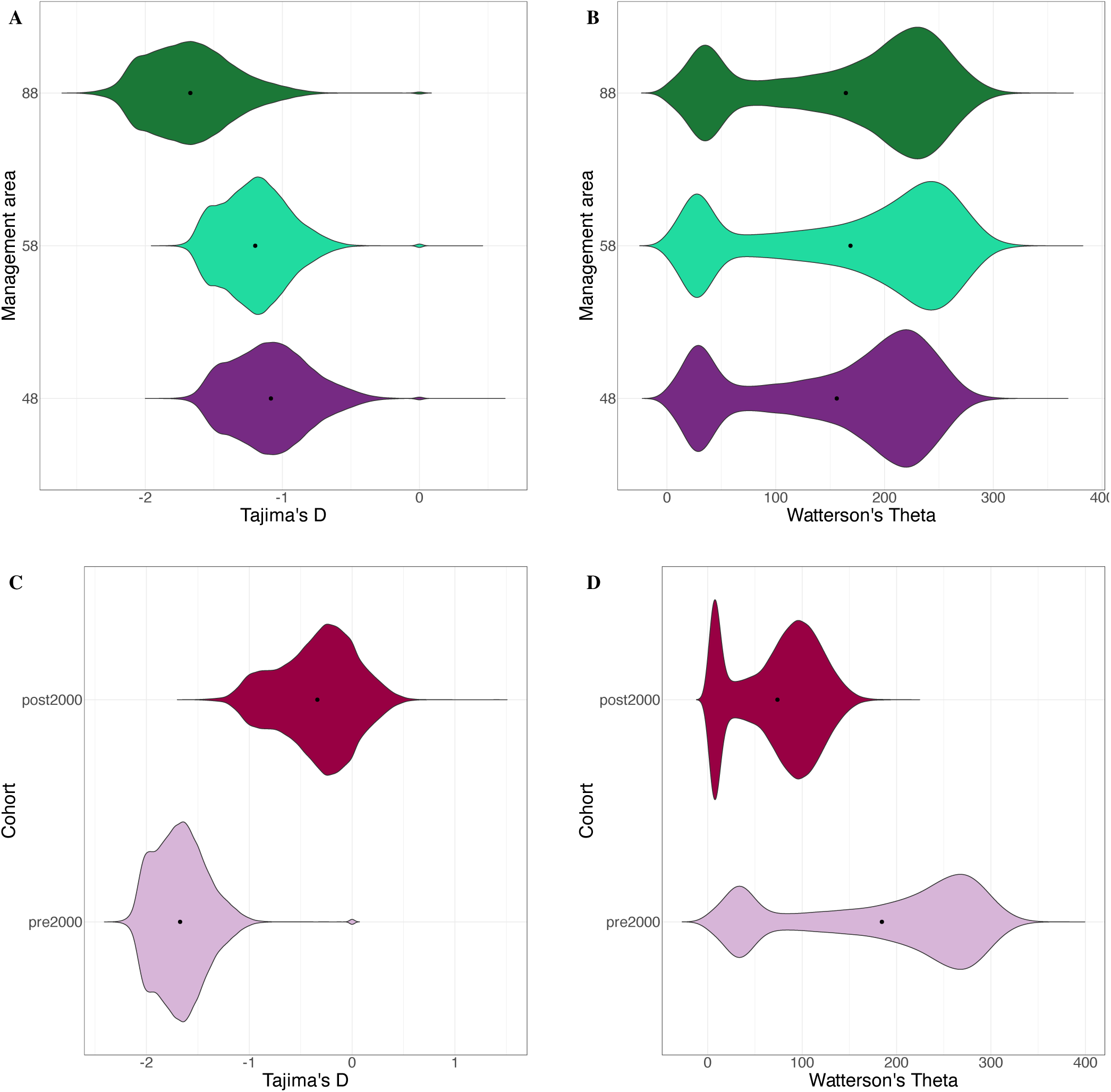
Violin plots of Tajima’s D and Watterson’s theta estimates derived from *D. mawsoni* genotype-likelihoods (2× dataset) using a sliding window approach with a window size of 50 kb and a step size of 10 kb; black dots indicate the mean value across the 50 kb-windows. (A) Tajima’s D and (B) Watterson’s theta estimates across CCAMLR management areas (*n* = 37). (C) Tajima’s D and (D) Watterson’s theta estimates between cohorts within area 88 (*n* = 18).

**Figure S10.**
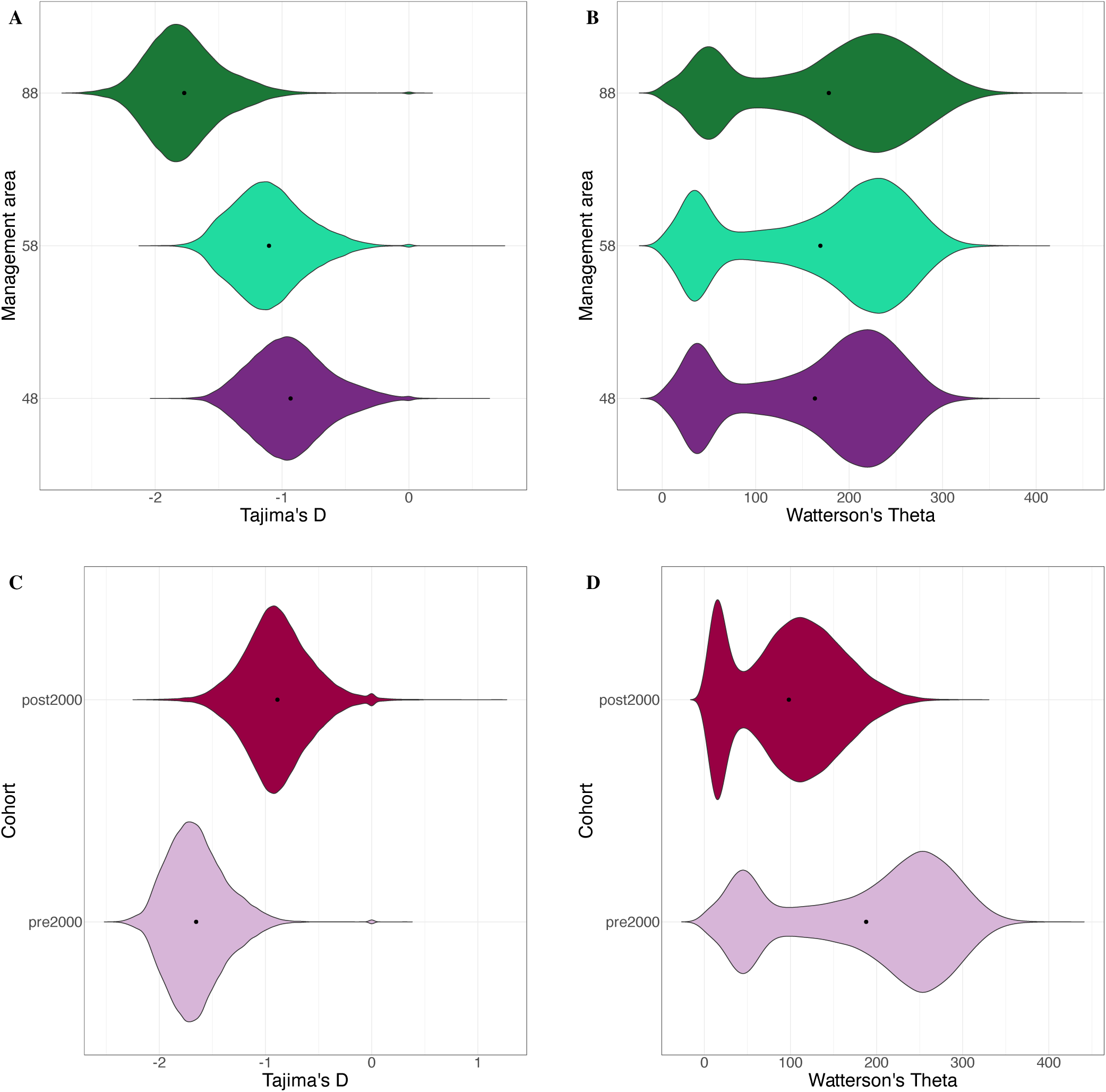
Violin plots of Tajima’s D and Watterson’s theta estimates derived from *D. mawsoni* genotype-likelihoods (noDS dataset) using a sliding window approach with a window size of 50 kb and a step size of 10 kb; black dots indicate the mean value across the 50 kb-windows. (A) Tajima’s D and (B) Watterson’s theta estimates across CCAMLR management areas (*n* = 41). (C) Tajima’s D and (D) Watterson’s theta estimates between cohorts within area 88 (*n* = 20).

